# The COX2-PGE2-PKA Axis Suppresses Antiviral Immunity by Inhibiting mtDNA-Dependent STING Activation

**DOI:** 10.64898/2026.04.03.716411

**Authors:** Pham Thuy Tien Vo, Julien Cicero, Zichen Wang, Hiroyuki Hakozaki, Thomas S Hoang, Sendi Rafael Adame-Garcia, Dana J. Ramms, Kuniaki Sato, Erica Stevenson, Yuan Zhou, Katie Fan, Danielle L. Swaney, Nevan J. Krogan, Johannes Schöneberg, Uri Manor, J. Silvio Gutkind

## Abstract

The innate immune cGAS-STING pathway is activated by cytosolic double-stranded DNA (dsDNA) to induce type I interferon (IFN) response, which is essential for mounting the antiviral response. However, STING activation during viral infection is often insufficient to achieve complete viral clearance, suggesting the existence of additional mechanisms that evade its activity. Here, we identified COX2/PGE_2_ as a negative regulator of STING activation, particularly in response to arising cytosolic mitochondrial DNA (mtDNA) generated during HSV-1 infection. Mechanistically, PGE_2_, through the EP4-cAMP-PKA axis, induces mitophagy to remove defective mitochondria and hence prevent the accumulation of immunostimulatory cytosolic mtDNA, thereby dampening STING-mediated type I IFN and antiviral response. Furthermore, we identified STOML2 as a downstream target of PKA that connects mitochondrial quality control with the regulation of innate immune signaling. Together, our findings establish the COX2/PGE_2_/PKA axis as a negative regulator of mtDNA-STING signaling that may be targeted to potentiate STING-mediated type I IFN and innate immunity.

Graphical abstract

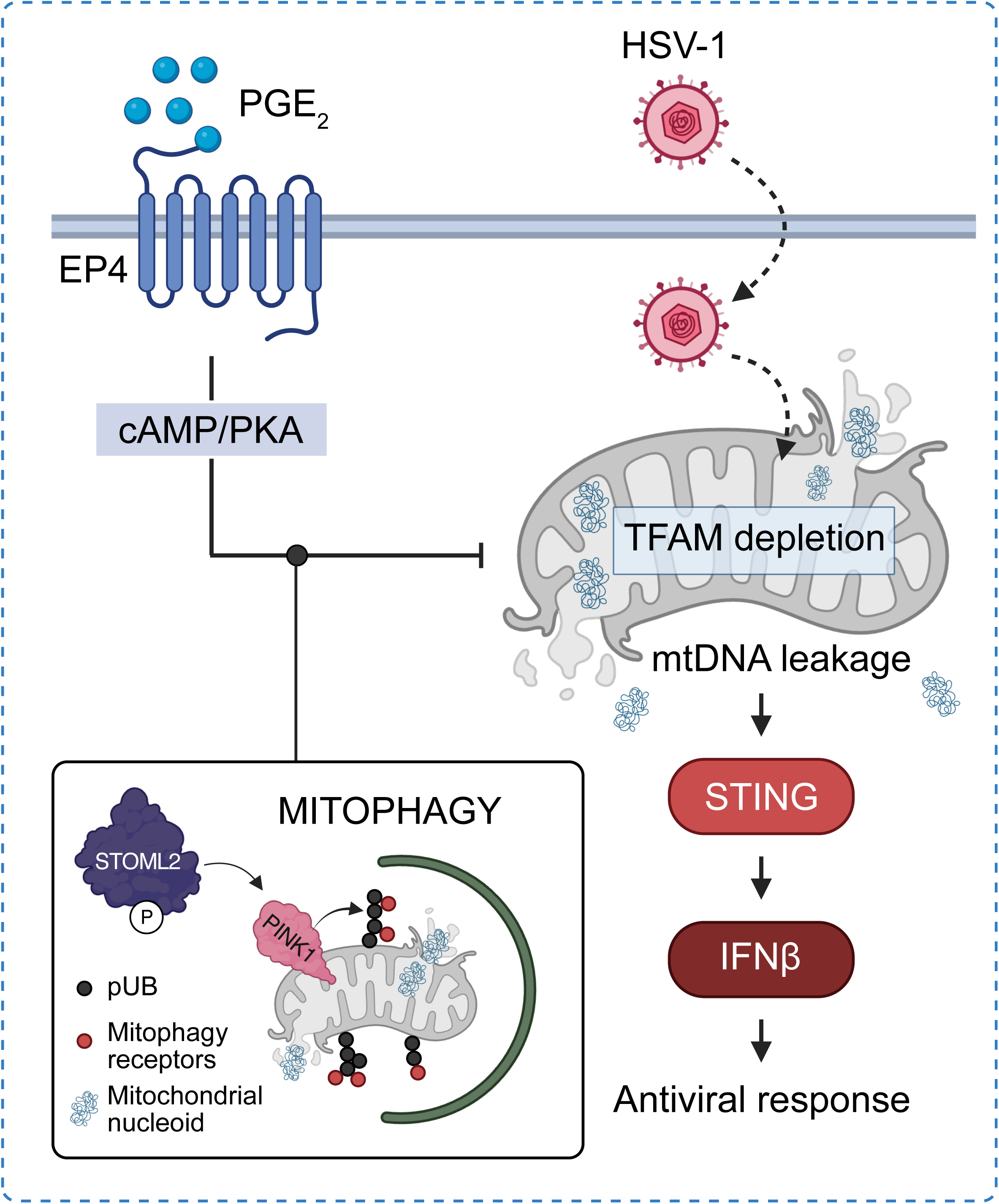

## INTRODUCTION

The human body must rapidly detect and respond to diverse threats arising from both external and internal sources, including pathogen infections, tissue damage, and cancer development. Innate immunity enables this surveillance by sensing molecular danger signals. Among these, the cyclic GMP-AMP synthase (cGAS)-stimulator of interferon genes (STING) pathway has emerged as a central signaling mechanism that detects cytosolic double-stranded DNA (dsDNA) derived from pathogens and from host sources such as nuclear or mitochondrial DNA^1–3^. Upon sensing cytosolic dsDNA, cGAS becomes activated and promotes the accumulation of the second messenger 2′3′-cyclic GMP–AMP (cGAMP). cGAMP subsequently binds to and activates STING, triggering a downstream signaling cascade that drives type I interferon (IFN) production and broad inflammatory gene expression programs that are essential for antiviral and anticancer immunity^2,4–8^. However, STING signaling activation during viral infection is often insufficient to achieve complete viral clearance^9,10^. This suggests the existence of yet-to-be-identified mechanisms that may counteract STING signaling. Given the need to potentiate STING signaling to enhance innate immunity, elucidating how viruses negatively regulate this pathway may reveal druggable molecular switches that can be leveraged to enhance STING activity in a context-dependent manner with the ultimate goal of improving patient outcomes.

Prostaglandin E2 (PGE_2_) is a prominent bioactive lipid mediator generated from arachidonic acid by the cyclooxygenase (COX) isoforms COX1 and COX2, with COX2 frequently upregulated in many inflammatory conditions^11,12^. PGE_2_ signals through G protein-coupled receptors (GPCRs) expressed on multiple immune cell populations, acting primarily via the Gαs-coupled receptors EP2 (PTGER2) and EP4 (PTGER4)^13^. Reports from our and other laboratories demonstrated that PGE_2_ promotes recruitment of myeloid-derived suppressor cells (MDSCs), reduces natural killer (NK) cell infiltration, and suppresses CD8^+^ T cell activation, thereby fostering an immunosuppressive tumor microenvironment^14–16^. While prior studies have largely focused on the role of PGE_2_ on adaptive immunity, whether and how PGE_2_ interferes with innate immunity, with STING signaling being a core component, during viral infection remains unclear.

Here, we addressed this knowledge gap and discovered that the COX2/PGE_2_/PKA axis forms a regulatory feedback loop and functions as a negative regulator of STING-mediated antiviral innate immunity, particularly in response to cytosolic mitochondrial DNA (mtDNA) generated during HSV-1 infection. Mechanistically, we demonstrated that PGE_2_ promotes mitophagy to eliminate defective mitochondria and hence curtail immunostimulatory cytosolic mtDNA, thereby dampening the downstream STING-mediated type I IFN and antiviral response in a EP4- and PKA-dependent manner. Specifically, we demonstrated that PKA induces PINK1-mediated mitophagy by phosphorylating STOML2 at Ser29 in response to PGE_2_ stimulation. Taken together, our findings suggest that COX2/PGE_2_/PKA may represent a regulatory node that can be targeted to enhance STING-mediated innate immunity.

## RESULTS

### PGE2 dampens antiviral response to HSV-1

To investigate whether PGE_2_ affects the antiviral response, we initially focused on whether this inflammatory mediator impacts herpes simplex virus 1 (HSV-1) infection, as the potent antiviral properties of STING signaling have been well characterized in the context HSV-1^17^ **(Fig. 1A)**. For this, murine bone marrow-derived macrophages (BMDMs) were infected with GFP-expressing HSV-1 for 24 hours in the presence or absence of PGE_2_ **(Fig. S1A)**. Whereas vehicle-treated macrophages exhibited resistance to HSV-1 infection, PGE_2_-treated macrophages displayed robust viral GFP expression post-infection **(Fig. 1B)**. Quantification of viral genome further revealed an approximately 2-fold increase in viral genomic abundance upon PGE_2_ stimulation **(Fig. 1C),** indicating that PGE_2_ dampens the antiviral response and hence promotes viral replication. Intriguingly, pharmacological inhibition of cyclooxygenase-2 (COX2), the enzyme responsible for PGE_2_ production **(Fig. 1D)**, reduced viral replication by approximately 50% **(Fig. 1E)**. Together, these findings suggest that the COX2/PGE_2_ axis may serve as a negative regulator of innate immune activation, thereby limiting antiviral response to HSV-1 and promoting viral replication.

**Figure 1.**
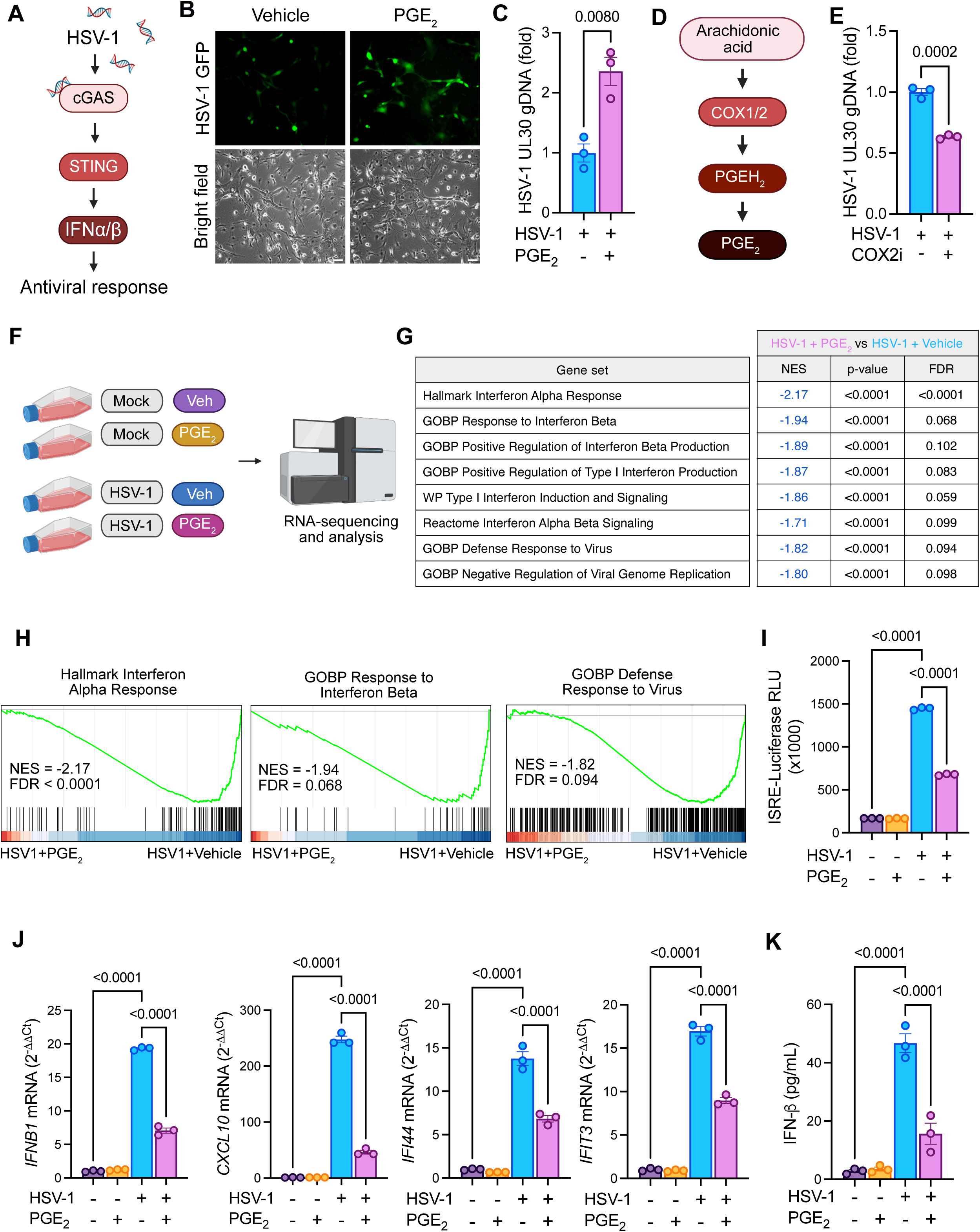
PGE_2_ hinders antiviral response to HSV-1. **(A)** Schematic of the antiviral property of cGAS/STING-mediated type I IFN during HSV-1 infection. **(B)** Representative images of bone marrow-derived macrophages infected with GFP-expressing HSV-1 (MOI = 1) for 24 hours in the presence or absence of 1 µM PGE_2_. **(C)** Bone marrow-derived macrophages were infected with HSV-1 (MOI = 1) in the presence or absence of 1 µM PGE_2_. At 24 h.p.i, cell lysates were collected and subjected to qPCR to quantify HSV-1 UL30 genomic abundance (n = 3). Data are shown as ΔΔCt values. **(D)** Schematic of the COX2/PGE_2_ signaling axis. **(E)** Bone marrow-derived macrophages were infected with HSV-1 (MOI = 1) in the presence or absence of 1 µM celecoxib. At 24 h.p.i, cell lysates were collected and subjected to qPCR to quantify HSV-1 UL30 genomic abundance (n = 3). Data are shown as ΔΔCt values. **(F)** Schematic illustrates experimental design of bulk RNA-seq workflow in THP-1 macrophages under four conditions: vehicle (Veh), PGE_2_ alone (PGE_2_), infected with HSV-1 for 16 hours in the absence (HSV-1+Veh) or presence of exogenous PGE_2_ (HSV-1+PGE_2_) to study mechanisms in which PGE_2_ controls innate immunity in response to HSV-1 infection. **(G-H)** Gene set enrichment analysis (GSEA) of RNA-seq data comparing HSV-1 + PGE2 versus HSV-1 + Veh. **(G)** Tables summarize enriched gene sets and associated statistics. **(H)** Representative enrichment plots for the indicated pathways. See also Fig. S1E. NES, normalized enrichment score; FDR, false discovery rate. **(I)** THP-1 macrophages expressing ISRE-Luciferase were mock-infected and infected with HSV-1 (MOI = 1) in the presence or absence of 1 µM PGE_2_. At 16 h.p.i, cell lysates were collected and subjected to a luciferase reporter assay to assess ISRE promoter activity (n = 3). **(J)** THP-1 macrophages were mock-infected and infected with HSV-1 (MOI = 1) in the presence or absence of 1 µM PGE_2_. At 16 h.p.i, cell lysates were collected and subjected to RT-qPCR to measure mRNA levels of representative type I ISGs (n = 3). Data are shown ΔΔCt values. **(K)** THP-1 macrophages were mock-infected and infected with HSV-1 (MOI = 1) in the presence or absence of 1 µM PGE_2_. At 24 h.p.i, supernatants were collected and analyzed by ELISA to measure secreted levels of IFNβ (n = 3). All experiments were performed with three independent biological replicates and repeated at least twice with reproducible results. Data are presented as mean ± s.e.m. Statistical significance was determined by unpaired, two-tailed Student’s *t-test* (C, E) or one-way ANOVA followed by Sidak’s multiple comparisons test (I-K); *p-*values are indicated.

To elucidate the mechanisms by which PGE_2_ controls innate immunity, we performed bulk RNA sequencing (RNA-seq) of THP-1-derived human macrophages that were treated with vehicle (Veh), PGE_2_ only (PGE_2_), or infected with HSV-1 for 16 hours in the absence (HSV-1+Veh) or presence of exogenous PGE_2_ (HSV-1+PGE_2_) **(Fig. 1F)**. Principal component analyses indicated clear separation between HSV-1+Veh and HSV-1+PGE_2_ samples, suggesting that PGE_2_ profoundly reshapes the transcriptional landscape triggered by HSV-1 infection **(Fig. S1B)**. Remarkably, gene set enrichment analysis (GSEA) comparing these two groups showed that gene sets involved in positive regulation of type I interferon response, including the interferon alpha and beta response, and defense response to virus infection, were among the most significantly downregulated pathways in macrophages treated with PGE_2_ **(Fig. 1G-H, S1C)**. It is noteworthy that more than 50% of genes downregulated by PGE_2_ stimulation were type I interferon-stimulated genes (ISGs) **(Fig. S1D-E)**. In line with this, transcription factor (TF) activity prediction analysis revealed that PGE_2_ led to a strong suppression of virus-induced TFs that control a broad range of type I ISGs expression, including STAT2 and IRF9 (**Fig. S1F-G**).

To validate the effect of PGE_2_ stimulation on virus-induced type I IFN, we measured the expression of representative type I ISGs that harbor IFN-sensitive response elements (ISREs) in their regulatory promoters, including *IFN1B*, *CXCL10*, *IFIT3*, and *IFI44.* Consistent with our RNA-seq analysis, HSV-1 infection robustly induced the activation of a reporter system driven by the ISRE promoter **(Fig. 1I)** and the transcription of type I ISGs; however, these responses were remarkably diminished by 50-70% in the presence of PGE_2_ in both THP-1-derived human macrophages **(Fig. 1J)** and murine BMDMs **(Fig. S1H)**. Moreover, PGE_2_ stimulation significantly suppressed virus-induced IFNβ secretion **(Fig. 1K)**, further establishing that PGE_2_ acts as a potent suppressor of type I IFN triggered by HSV-1 infection.

### HSV-1 infection induces the leakage of immunostimulatory mtDNA into the cytosol that contributes to STING-mediated antiviral innate immunity

While the type I IFN response can be activated by other innate immune pathways in addition to STING activation, our GSEA analysis specifically identified gene sets implicated in endogenous cytosolic DNA sensing as the most enriched pathways in macrophages challenged with HSV-1, further confirming that STING signaling is the prominent driver of antiviral response in our model **(Fig. S2A)**. This prompted us to assess the effect of PGE_2_ on the activation of the STING signaling pathway following HSV-1 infection **(Fig. 2A)**. Our data revealed that viral infection markedly increased phosphorylation of TBK1 and IRF3; however, both were strongly suppressed upon PGE_2_ stimulation **(Fig. 2B-D)**. Based on this result, we next asked whether PGE_2_ directly affects STING activation. Since PGE_2_ had no effect on ISRE promoter activation elicited by the STING agonist 2’3’-cGAMP **(Fig. S2B)**, we therefore concluded that PGE_2_ functions upstream of STING activation.

**Figure 2.**
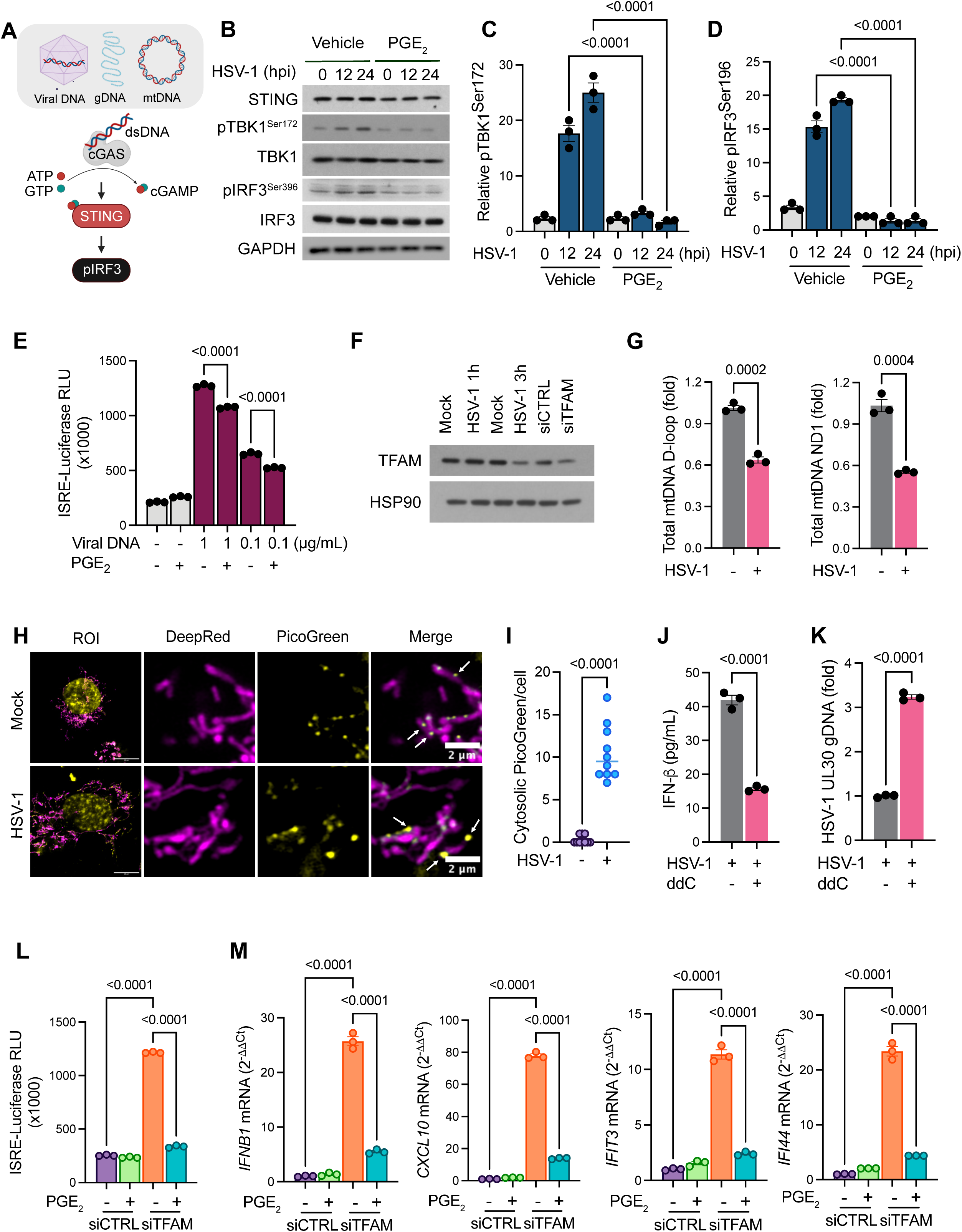
PGE_2_ exerts inhibitory effects on STING-mediated innate immunity arising from immunostimulatory cytosolic mtDNA generated during viral infection. **(A)** A schematic illustrates the activation of STING, TBK1, and IRF3 in response to double-stranded DNA (dsDNA) in the cytosol by cGAS. **(B-D)** THP-1 macrophages were mock-infected or infected with HSV-1 in the presence or absence of 1 µM PGE_2_. At the indicated h.p.i, cell lysates were collected and subjected to immunoblotting with the indicated antibodies **(B)**. Band intensity of phosphorylated TBK1 **(C)** and IRF3 **(D)** was quantified and normalized to total TBK1 and IRF3, respectively (n = 3). **(E)** THP-1 macrophages expressing ISRE-Luciferase were transfected with viral DNA at indicated concentrations in the presence or absence of 1 µM PGE_2_ for 16 hours. Cell lysates were collected and subjected to a luciferase reporter assay to assess ISRE promoter activity (n = 3). **(F)** THP-1 macrophages were mock-infected or infected with HSV-1 (MOI = 1) for the indicated times. Cell lysates were collected and subjected to immunoblotting with the indicated antibodies. Samples transfected with either scramble siRNA (siCTRL) or TFAM siRNA (siTFAM) were included as controls. **(G)** THP-1 macrophages were mock-infected or infected with HSV-1 (MOI = 1). At 16 h.p.i, cell lysates were collected and subjected to qPCR to assess levels of total mt-D-loop and mt-ND1 regions (n = 3). **(H)** Representative Airyscan live-cell imaging of THP-1 macrophages mock-infected or infected with HSV-1 for 3 hours. MitoTracker DeepRed (magenta): mitochondria, PicoGreen (yellow): DNA. Scale bars, 2 µm. **(I)** Quantification of cytosolic PicoGreen in THP-1 macrophages mock-infected or infected with HSV-1 (MOI = 1) for 3 hours (n = 10). **(J-K)** THP-1 macrophages were infected with HSV-1 in the presence or absence of the mtDNA replication inhibitor, ddC. At 24 h.p.i, cell supernatants were collected and subjected to ELISA to measure levels of secreted IFNβ (n = 3) **(J)**, and cell lysates were collected and subjected to qPCR to quantify HSV-1 UL30 genomic abundance (n = 3) **(K)**. **(L)** THP-1 macrophages expressing ISRE-Luciferase were transfected with either scramble siRNA (siCTRL) or TFAM siRNA (siTFAM) in the presence or absence of 1 µM PGE_2._ At 16 hours post stimulation, cell lysates were collected and subjected to a luciferase reporter assay to assess ISRE promoter activity (n = 3). **(M)** THP-1 macrophages were transfected with either scramble siRNA (siCTRL) or TFAM siRNA (siTFAM) in the presence or absence of 1 µM PGE_2_. At 16 hours post-stimulation, cell lysates were collected and subjected to RT-qPCR to assess the mRNA levels of representative type I ISGs (n = 3). All experiments were performed with three independent biological replicates and repeated at least twice with reproducible results. Data are presented as mean ± s.e.m. Statistical significance was determined by one-way ANOVA followed by Sidak’s multiple comparisons test (**C-E, L-M**) or unpaired, two-tailed Student’s t-test (**G, I-K**). *p*-values are indicated.

Given that cGAS can be activated by both viral and host-derived DNA^1,3,18^, we next assessed whether the inhibitory effect of PGE_2_ can be recapitulated using viral DNA as the stimulus. To our surprise, although PGE_2_ significantly attenuated viral DNA-induced ISRE activation **(Fig. 2E)**, the magnitude of inhibition was substantially lower than that observed during HSV-1 infection **(Fig. S2C)**. While we do not exclude the inhibitory effect of PGE_2_ on viral DNA-induced innate immunity, this discrepancy led us to consider the possibility that PGE_2_ may control type I IFN elicited by non-viral DNA sources generated during viral infection.

In addition to viral DNA, cGAS can also detect host-derived DNA, including nuclear and mitochondrial DNA^3,18^, in the cytosol. Intriguingly, previous studies have demonstrated that an HSV-1-encoded protein, UL12.5, targets and depletes the mitochondrial transcription factor A (TFAM), which controls mtDNA replication and nucleoid organization^19–21^. This results in the reduction in total mtDNA number and subsequent release of mtDNA into the cytosol, where it activates the STING innate immune signaling^3^. Indeed, we observed that HSV-1 infection reduced both TFAM protein abundance **(Fig. 2F)** and total mtDNA copy number **(Fig. 2G)** as early as 3 hours post-infection in our macrophage model. To determine whether HSV-1 infection induces cytosolic mtDNA release, we performed live-cell imaging with PicoGreen to label DNA and MitoTracker DeepRed to label mitochondria in mock- and HSV-1-infected macrophages. Airyscan imaging revealed a subset of enlarged nucleoids located outside the mitochondria in infected cells, a phenomenon that was not observed in the control **(Fig. 2H-I)**, indicating that HSV-1 infection indeed leads to the release of mtDNA into the cytosol.

To determine whether mtDNA is required for STING-mediated type I IFN induction and antiviral response, we treated HSV-1-infected macrophages with dideoxycytidine (ddC), a deoxyribonucleoside analog that selectively inhibits mtDNA replication and reduces mtDNA copy number^22^ **(Fig. S2D)**. Remarkably, the treatment of ddC significantly attenuated type I IFN, as indicated by reduced IFNβ secretion **(Fig. 2J)**, suppressed ISRE promoter activation **(Fig. S2E)**, and resulted in a 3-fold increase in viral genome abundance post-infection **(Fig. 2K)**. Together, these data demonstrate that HSV-1 infection promotes the release of mtDNA into the cytosol and that, in turn, cytosolic mtDNA is necessary to fully engage STING-mediated type I IFN and antiviral innate immunity.

### PGE_2_ exerts inhibitory effects on STING-mediated innate immunity arising from immunostimulatory cytosolic mtDNA generated during viral infection

Having confirmed mtDNA as a critical endogenous trigger during HSV-1 infection, we next asked whether PGE_2_ controls mtDNA-dependent activation of STING signaling. To model mtDNA release independently of viral infection, macrophages were transfected with TFAM small interfering RNA (siRNA). Consistent with prior reports^3^, TFAM depletion robustly induced ISRE promoter activation in a STING-dependent manner **(Fig. S2F)**. Strikingly, we found that TFAM siRNA-induced innate immune responses were suppressed by 70-80% in the presence of PGE_2_ stimulation **(Fig. 2L-M)**, supporting that PGE_2_ dampens virus-induced innate immunity, at least in part, by inhibiting mtDNA-dependent activation of STING signaling.

To determine whether the inhibitory effect of PGE_2_ on the mtDNA-STING signaling axis is restricted to macrophages, we extended our analysis to HEK293 cells co-expressing cGAS-STING and PGE_2_ receptor, EP4. Consistent with our observations in human macrophages, PGE_2_ stimulation remarkably suppressed the induction of type I ISGs triggered by TFAM depletion in this cell line (**Fig. S3A-B**), confirming that the effect of PGE_2_ on mtDNA-STING signaling is a broad biological mechanism rather than cell-type specific.

### COX2/PGE_2_ is upregulated in response to mtDNA-dependent STING activation

Given that PGE_2_ serves as a negative regulator of mtDNA-STING-mediated innate immunity, we next asked whether PGE_2_ and its upstream enzyme COX2 are induced upon mtDNA-STING activation, thereby forming a novel feedback regulatory loop. To test this hypothesis, we examined COX2 transcript levels in physiological contexts in which mtDNA acts as an endogenous agonist to activate STING signaling. Intriguingly, we observed a marked increase in COX2 transcript levels in mice treated with doxorubicin, a first-line chemotherapeutic agent that was reported to induce mtDNA-STING signaling^23^ **(Fig. 3A)**. Similarly, analysis of a separate dataset on proliferating and senescent cells^24^ revealed elevated COX2 expression in senescent cells, which exhibited mtDNA-STING activation **(Fig. 3B)**. It is noteworthy that we also detected a significant increase in both COX2 transcript levels **(Fig. 3C)** and COX2-induced extracellular PGE_2_ release **(Fig. 3D)** following HSV-1 infection at 24 and 48 hours post infection in our macrophage model, further supporting the conclusion that COX2/PGE_2_ is upregulated downstream of mtDNA-STING activation to establish a feedback loop that restrains STING activation.

**Figure 3.**
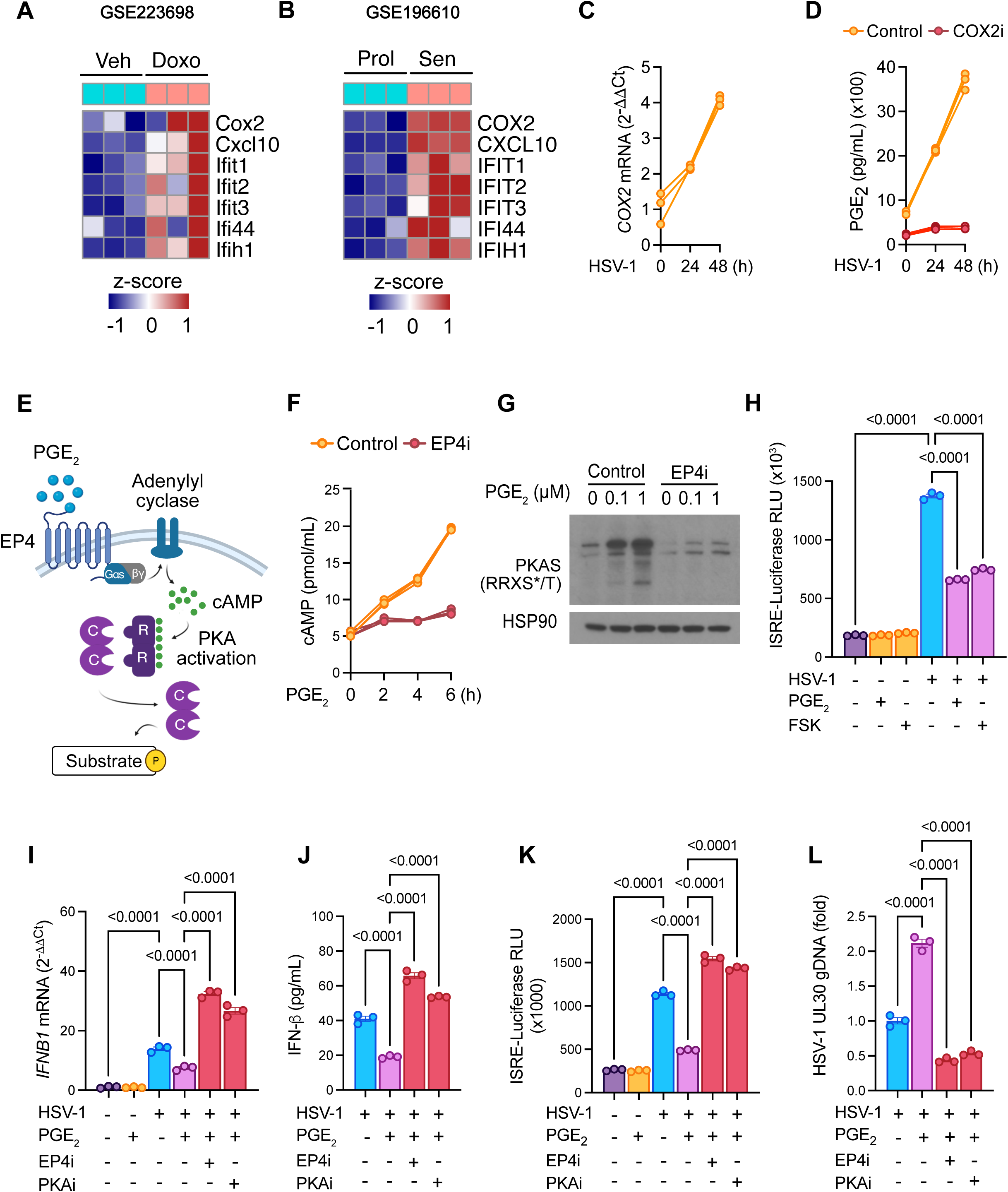
PGE_2_ signals via EP4-cAMP-PKA axis to function as a negative regulator of mtDNA-STING-mediated type I IFN and antiviral innate immunity. **(A)** Heatmap showing relative expression of COX2 and type I ISGs in mice treated with either vehicle (Veh) or doxorubicin (Doxo) (GEO: GSE223698). **(B)** Heatmap showing relative expression of COX2 and type I ISGs in proliferating cells (Prof) and senescence cells (Sen) (GEO: GSE196610). **(C)** THP-1 macrophages were mock-infected or infected with HSV-1 (MOI = 1) for either 24 or 48 hours. Cell lysates were subjected to RT-qPCR to assess mRNA levels of COX2 (n = 3). **(D)** THP-1 macrophages were mock-infected or infected with HSV-1 (MOI = 1) for either 24 or 48 hours in the presence or absence of 1 µM celecoxib (COX2i). Cell lysates were collected and analyzed by ELISA to measure extracellular PGE_2_ levels (n = 3). **(E)** A schematic illustrates PGE_2_-cAMP-PKA signaling. **(F)** THP-1 macrophages were treated with 1 µM PGE_2_ in the presence or absence of 1 µM EP4 inhibitor (EP4i) for 16 hours. Cell lysates were collected and analyzed by ELISA to measure intracellular cAMP levels (n = 3). **(G)** THP-1 macrophages were treated with 1 µM PGE_2_ at indicated concentrations in the presence or absence of 1 µM EP4 inhibitor (EP4i) for 16 hours. Cell lysates were collected and subjected to immunoblotting with the indicated antibodies. **(H)** THP-1 macrophages expressing ISRE-Luciferase were mock-infected or infected with HSV-1 (MOI = 1) in the presence of either 1 µM PGE_2_ or 1 µM forskolin and 50 µM IBMX. At 16 h.p.i, cell lysates were collected and subjected to a luciferase reporter assay to assess ISRE promoter activity (n = 3). **(I-J)** THP-1 macrophages were mock-infected or infected with HSV-1 (MOI = 1), followed by 1 µM PGE_2_ stimulation in the presence of either 1 µM EP4 inhibitor (EP4i) or 1 µM PKA inhibitor (PKAi). At 24 h.p.i, cell lysates were subjected to RT-qPCR to assess mRNA levels of IFNβ (n = 3) **(I)**, and cell supernatants were subjected to ELISA to measure secreted levels of IFNβ (n = 3) **(J)**. **(K)** THP-1 macrophages expressing ISRE-Luciferase were mock-infected or infected with HSV-1 (MOI = 1), followed by 1 µM PGE_2_ stimulation in the presence of either 1 µM EP4 inhibitor (EP4i) or 1 µM PKA inhibitor (PKAi). At 16 h.p.i, cell lysates were collected and subjected to a luciferase reporter assay to assess ISRE promoter activity (n = 3). **(L)** THP-1 macrophages were infected with HSV-1 (MOI = 1), followed by 1 µM PGE_2_ stimulation in the presence of either 1 µM EP4 inhibitor (EP4i) or 1 µM PKA inhibitor (PKAi). At 16 h.p.i, cell lysates were collected and subjected to RT-qPCR to assess HSV-1 UL30 genomic abundance (n = 3). Data are presented as mean ± s.e.m. Statistical significance was determined by one-way ANOVA followed by Sidak’s multiple comparisons test. *P*-values are indicated.

### PGE_2_ signals through EP4-cAMP-PKA axis to function as a negative regulator of mtDNA-STING-mediated antiviral innate immunity

To elucidate the downstream mechanism, we next sought to determine the signaling cascade that facilitates the intracellular effects of PGE_2_. As anticipated, PGE_2_ stimulation triggered rapid and sustained accumulation of the second messenger cyclic adenosine monophosphate (cAMP), accompanied by activation of its downstream effector, protein kinase A (PKA), in macrophages **(Fig. 3E-G)**^25^. Indeed, direct activation of cAMP signaling with forskolin and IBMX stimulation recapitulated the suppressive effect of PGE_2_ on ISRE promoter activation (**Fig. 3H)**. Given that PGE_2_ signals through G protein-coupled receptors, EP1-EP4, and that we did not detect expressions of EP1, EP2, or EP3 receptors in our model^26^, we reasoned that the observed effects of PGE_2_ are mediated predominantly through EP4. Indeed, pharmacological inhibition of EP4 receptors significantly abrogated the increase in intracellular cAMP and PKA activation induced by PGE_2_ **(Fig 3F-G)**. Furthermore, inhibition of either EP4 or PKA, the latter with a first-generation PKA-selective inhibitor^27^, not only fully reversed the inhibitory effect of PGE_2_ on STING-mediated type I IFN but also enhanced antiviral response, as indicated by elevated IFNβ expression **(Fig. 3I-J)**, heightened ISRE promoter activation **(Fig. 3K)**, and reduced viral genome abundance **(Fig. 3L)** following HSV-1 infection. Based on these findings, we concluded that PGE_2_ signals via the EP4-cAMP-PKA axis to function as a negative regulator of mtDNA-STING signaling.

### Protein-protein interaction network analysis reveals a novel link between PKA Cα and the mitochondrial quality control machinery

To further investigate the interplay between PGE_2_-PKA signaling and mtDNA-STING pathway, we sought to identify downstream targets of PKA that may mediate the effects of PGE_2_ on innate immunity regulation. PKA functions as a holoenzyme that consists of two regulatory subunits and two catalytic subunits. Once activated, cAMP binds to the regulatory subunit and releases the active catalytic subunits to phosphorylate target proteins^28–31^. Among the isoforms, PKA catalytic subunit alpha (PKA Cα) is the most commonly expressed across human tissues^30^.

For our purpose, we generated a panel of HEK293 cell lines harboring doxycycline-inducible expression of either PKA Cα wild-type or a W197R mutant that exhibits reduced affinity for the regulatory subunit, both of which are fused to a C-terminal Flag tag **(Fig. S4A-B)**. Given that a complete disengagement of the regulatory subunit can enhance binding of certain interactors to PKA Cα^32^, the inclusion of the W197R mutant enabled us to capture a comprehensive spectrum of the PKA Cα interactome. Cells were treated with or without doxycycline for 48 hours, followed by anti-FLAG immunoprecipitation and affinity purification-mass spectrometry (AP-MS) **(Fig. 4A)**. Consistent with previous reports showing implications of PKA activity in a broad range of physiological contexts^30^, our proteomic analysis revealed a diverse repertoire of candidate PKA Cα interactors across many cellular processes, ranging from metabolism, ribosomal biogenesis, to RNA-binding proteins and DNA replication **(Fig. S4C-D)**. Strikingly, by comparing the interactions detected in our PKA interactome with those documented in publicly available databases (e.g., BioPlex, STRING, CORUM, BioGRID, and IMEX), we found that over 86% of our interactions are novel and not previously reported **(Fig. S4E)**. Although canonical components of the STING pathway were not among the top interactors, our data revealed novel interactions between PKA Cα and multiple regulators of the mitochondrial quality control machinery. We next performed KEGG pathway analysis to gain insights into their biological functions and unexpectedly uncovered a significant enrichment for mitophagy pathway, which is defined as the removal of faulty or superfluous mitochondria by the autophagy machinery **(Fig. 4B).** Given that defective mitochondria represent a major source of cytosolic mtDNA, we therefore hypothesized that PGE_2_-PKA activation may promote mitophagy as a mechanism to eliminate damaged mitochondria. Such a mechanism would be predicted to limit cytosolic mtDNA release and thereby attenuate subsequent STING-mediated type I IFN and innate immune activation.

**Figure 4.**
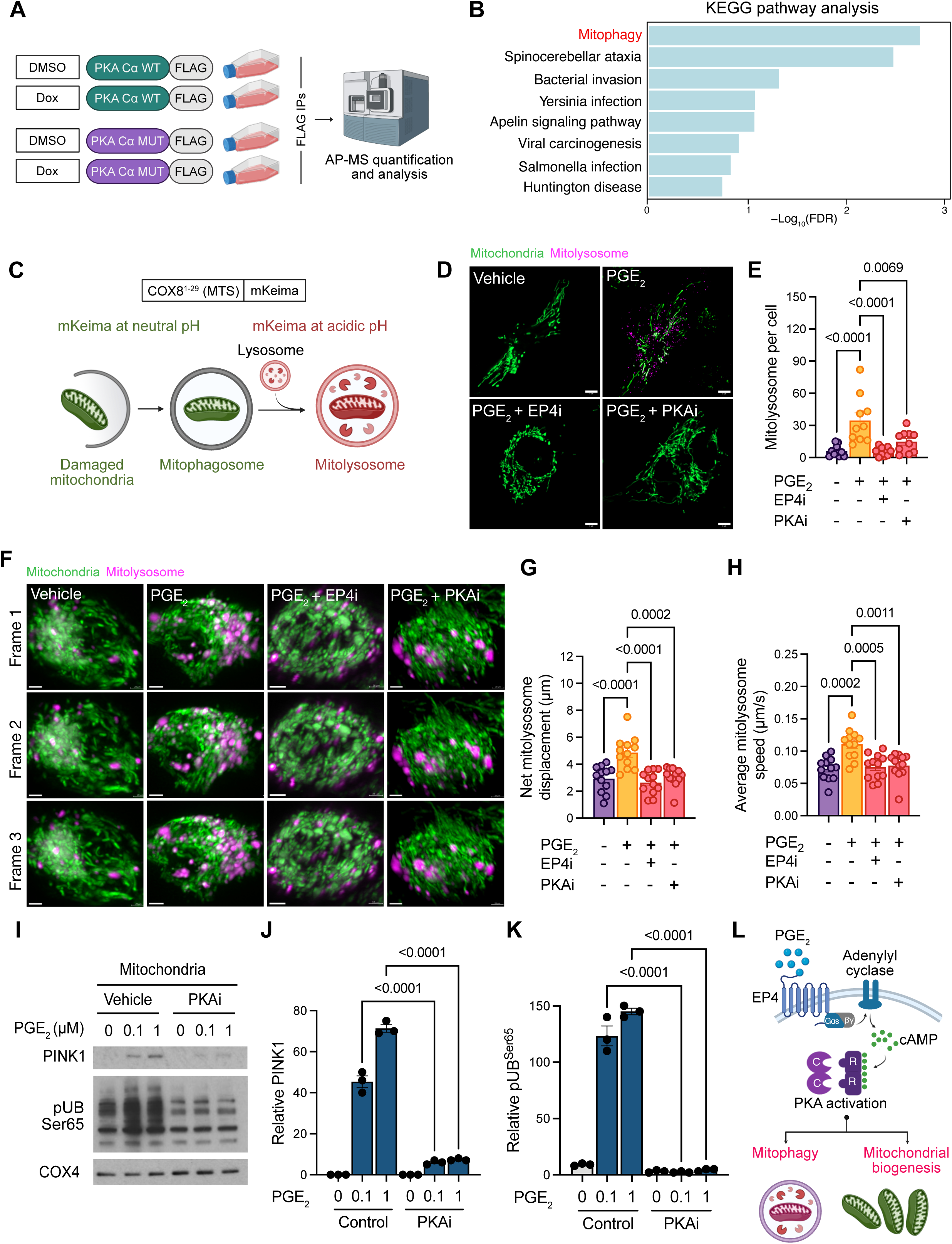
PGE_2_ stimulation promotes PINK1-mediated mitophagy in a PKA- and EP4-dependent manner. **(A)** Proteomic workflow in HEK293 cells with doxycycline-inducible expression of PKA Cα wild-type (WT) or W197R mutant (MUT). **(B)** The top 8 significantly enriched KEGG pathways within the mitochondrial quality control category identified from the proteomic dataset. **(C)** Schematic of the mt-mKeima mitophagy reporter. **(D)** Representative Airyscan live-cell imaging of THP-1 macrophages expressing mt-mKeima sensor treated with 1 µM PGE_2_ in the presence of either 1 µM EP4 inhibitor (EP4i) or 1 µM PKA inhibitor (PKAi) for 16 hours. Scale bar, 10 µm. **(E)** Quantification of mitolysosome numbers shown in **(D)** (n = 10). Data were quantified from one representative experiment of three. **(F)** Representative images of live-cell 4D lattice light sheet imaging on THP-1 macrophages treated as in **(D)**. Scale bar, 20 µm. **(G-H)** Quantification of net mitolysosome displacement (n = 10) **(G)** and average mitolysosome speed (n = 12) **(H)** from imaging in **(F)**. **(I)** THP-1 macrophages treated with PGE_2_ at indicated concentrations in the presence or absence of 1 µM PKA inhibitor (PKAi) for 16 hours. Mitochondrial fractions were isolated and subjected to immunoblotting with indicated antibodies. Band intensities were quantified and normalized to COX4 expression for PINK1 **(J)** and pUB Ser65 **(K)** (n = 3). **(L)** Working model illustrating that PGE_2_ induces mitophagy and mitochondrial biogenesis to enhance mitochondrial homeostasis in a EP4- and PKA-dependent manner. Data are presented as mean ± s.e.m. Statistical significance was determined by one-way ANOVA followed by Sidak’s multiple comparisons test. *p*-values are indicated.

### PGE_2_ stimulation promotes PINK1-mediated mitophagy in a PKA- and EP4-dependent manner

To test this hypothesis, we first validated the effect of the PGE_2_-PKA axis on mitophagy using the fluorescent protein, mKeima, fused to a mitochondrial targeting sequence (mtKeima). This sensor shifts its excitation wavelength in response to the pH of its environment. Given that during mitophagy, mitochondria move from the neutral pH environment of the cytoplasm (green) to the acidic environment of the lysosome (magenta), mtKeima enables us to monitor mitophagy in live cells^33^ **(Fig. 4C)**. Airyscan imaging of macrophages stably expressing mtKeima revealed that PGE_2_ stimulation robustly activated mitophagy, as evidenced by a striking increase in the number of mitolysosomes, in an EP4- and PKA-dependent manner **(Fig. 4D-E)**. In addition, by leveraging *state-of-the-art* 4D lattice light-sheet microscopy, we demonstrated for the first time that PGE_2_-EP4-PKA signaling enhances mitolysosome displacement and speed in macrophages **(Fig. 4F-H)**, further indicating that PGE_2_-PKA activation leads to an increase in mitophagy flux. Given that mitophagy can be mediated by multiple mechanisms, including the PTEN-induced kinase 1 (PINK1)/Parkin pathway, mitophagy receptors, and changes in mitochondrial lipid composition^34,35^, we next assessed the effect of PGE_2_ on these pathways. Indeed, there was an increase in PINK1 recruitment and its downstream phosphorylation on Ubiquitin at Ser65 in macrophages upon PGE_2_-PKA activation **(Fig. 4I-K)**.

Although mitophagy is generally considered beneficial for mitochondrial homeostasis, excessive mitophagy can become detrimental due to uncontrolled mitochondrial mass depletion^36^. While this is highly unlikely in our model, given that PGE_2_ stimulation for 24 hours resulted in only a slight decrease in total mitochondrial mass **(Fig. S5A)**, we nonetheless evaluated the impact of PGE_2_ on mitochondrial function. Mitochondrial respirometry analysis revealed that PGE_2_ did not impair oxygen consumption, but instead slightly increased spare respiratory capacity **(Fig. S5B-C)**. Moreover, lower levels of mitochondrial ROS production were also observed in macrophages stimulated with PGE_2_ for 24 hours **(Fig. S5D)**. Notably, our RNA-seq analysis further revealed that PGE_2_ stimulation led to an increase in the transcript levels of PGC-1α, a master regulator of mitochondrial biogenesis^37^ **(Fig. S5E)**. Together, these data suggest that PGE_2_ may simultaneously stimulate mitochondrial biogenesis after mitophagy induction to replenish the healthy mitochondrial pool, thus promoting mitochondrial quality control **(Fig. 4L).**

### PINK1-mediated mitophagy induction by PGE_2_ mediates the reduction in cytosolic mtDNA and subsequent inhibition of STING-mediated antiviral type I IFN following viral infection

Consistent with the findings above, HSV-1-infected macrophages stimulated with either PGE_2_ or forskolin/IBMX exhibited increased PINK1-induced phosphorylation of Ubiquitin on Ser65 **(Fig. 5A)**. Since mitophagy promotes the clearance of damaged mitochondria, we reasoned that PGE_2_, by promoting mitophagy induction, may limit the release of mtDNA into the cytosol. Indeed, Airyscan imaging revealed that PGE_2_ markedly abrogated TFAM depletion-induced cytosolic DNA, as indicated by a significant decrease in enlarged Picogreen-positive nucleoid localized outside of the mitochondria **(Fig. 5B-C)**. As an orthogonal approach to further validate this result, we directly quantified cytosolic mtDNA using quantitative PCR (qPCR) **(Fig. 5D)**. Consistent with our imaging results, analysis of pure cytosolic fractions revealed a greater than 2-fold reduction in mtDNA fragments corresponding the D-loop and ND1 region upon PGE_2_ stimulation in both TFAM-depleted **(Fig. 5E)** and HSV-1-infected macrophages **(Fig. 5F)**, further confirming the effect of PGE_2_ in limiting mtDNA release. Consistently, these effects are EP4- and PKA-dependent **(Fig. S6A)**.

**Figure 5.**
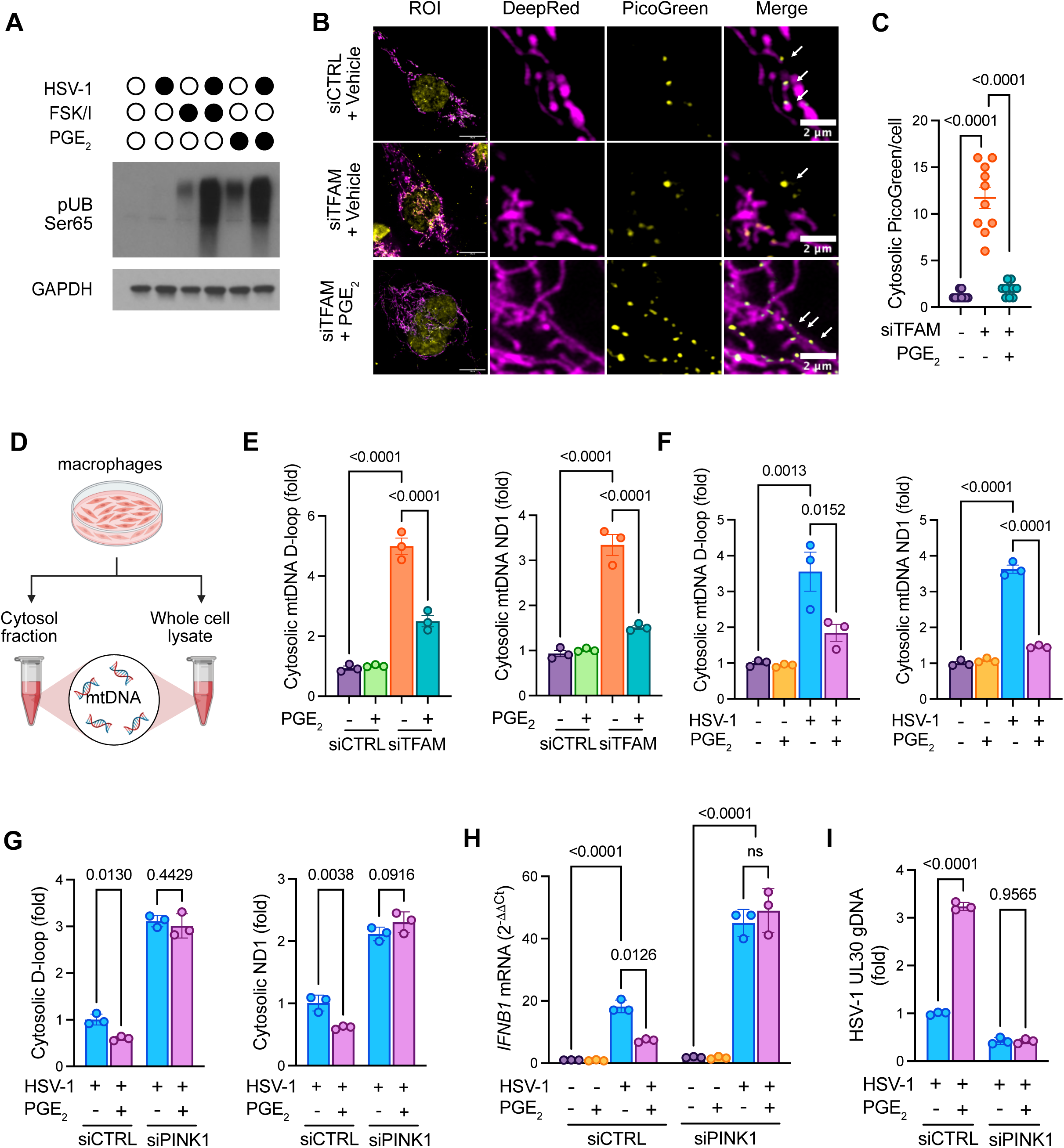
PINK1-mediated mitophagy induction by PGE_2_ mediates the reduction in cytosolic mtDNA and inhibition of STING-mediated antiviral type I IFN following viral infection. **(A)** THP-1 macrophages were treated with 1 µM PGE_2_ alone, 1 µM forskolin and 50 µM IBMX alone, or infected with HSV-1 (MOI = 1) in the presence or absence of PGE_2_ or forskolin/IBMX. At 16 h.p.i, cell lysates were collected and subjected to immunoblotting with indicated antibodies. **(B)** Representative Airyscan live-cell imaging of THP-1 macrophages transfected with either control siRNA (siCTRL) or TFAM siRNA (siRNA) in the absence or presence of 1 µM PGE_2_ for 16 hours. Scale bars, 2 µm. **(C)** Quantification of cytosolic PicoGreen from images in **(B)** (n = 10). Data were quantified from one representative experiment of three. **(D)** Schematic showing the workflow to quantify cytosolic mitochondrial DNA using qPCR. **(E)** THP-1 macrophages were transfected with either scramble siRNA (siCTRL) or TFAM siRNA (siRNA) in the presence or absence of 1 µM PGE_2_ for 16 hours. Cytosol fractions were isolated and subjected to qPCR to assess the presence of mt-Dloop and mt-ND1 regions (n = 3). **(F)** THP-1 macrophages were mock-infected or infected with HSV-1 (MOI = 1) in the presence or absence of 1 µM PGE_2_. At 16 h.p.i, cytosol fractions were isolated and subjected to qPCR to assess the presence of mt-Dloop and mt-ND1 regions (n = 3). **(G)** THP-1 macrophages were transfected with either scramble siRNA (siCTRL) or PINK1 siRNA (siPINK1), followed by mock or HSV-1 infection in the presence or absence of 1 µM PGE_2_. At 16 h.p.i, cytosol fractions were isolated and subjected to qPCR to assess the presence of mt-Dloop and mt-ND1 regions (n=3). **(H)** THP-1 macrophages were transfected with either scramble siRNA (siCTRL) or PINK1 siRNA (siPINK1), followed by mock or HSV-1 infection in the presence or absence of 1 µM PGE_2_. At 16 h.p.i, whole cell lysates were collected and subjected to RT-qPCR to assess mRNA levels of IFNβ (n = 3). **(I)** THP-1 macrophages were transfected with either scramble siRNA (siCTRL) or PINK1 siRNA (siPINK1), followed by mock or HSV-1 infection in the presence or absence of 1 µM PGE_2_. At 16 h.p.i, whole cell lysates were collected and subjected to qPCR to assess HSV-1 UL30 genomic abundance (n=3). All experiments were performed with three independent biological replicates and repeated at least twice with reproducible results. Data are presented as mean ± s.e.m. Statistical significance was determined by one-way ANOVA followed by Sidak’s multiple comparisons test. *p*-values are indicated.

To fully ascertain whether mitophagy is solely responsible for the PGE_2_-mediated inhibitory effect on cytosolic mtDNA, we evaluated the same readout for mtDNA release in macrophages in which PINK1-dependent mitophagy was downregulated **(Fig. S6B)**. Our data revealed that depletion of PINK1 fully reversed the PGE_2_-mediated decrease in cytosolic mtDNA D-loop and ND1 fragments in macrophages following HSV-1 viral infection **(Fig. 5G)**, indicating that this pathway is indeed responsible for curtailing mtDNA release. Furthermore, PINK1 depletion completely restored STING-mediated type I IFN and antiviral response that was otherwise suppressed by PGE_2_, as evidenced by elevated IFNβ expression **(Fig. 5H)** and reduced viral genome abundance **(Fig. 5I)**. Based on these results, we concluded that PGE_2_ induces PINK1-mediated mitophagy activation to improve mitochondrial quality control while limiting the cGAS upstream trigger, cytosolic mtDNA, thereby attenuating downstream STING-mediated type I IFN and antiviral response following HSV-1 infection.

### The expression of STOML2, a PKA interactor, is necessary for PINK1-mediated mitophagy induction and STING-mediated antiviral type I IFN inhibition upon PGE_2_ stimulation

Building on these intriguing findings, we revisited our PKA interactome to identify candidate PKA interactors that may mediate the effect of PGE_2_ on PINK1-dependent mitophagy, and the subsequent STING-mediated antiviral innate immunity following HSV-1 infection. Among the multiple PKA-interacting molecules, stomatin-like protein 2 (STOML2) emerged as a particularly compelling candidate **(Fig. 6A)**. Our proteomic analysis indicated that STOML2 preferentially interacts with wild-type PKA Cα, but not with the mutant PKA Cα, suggesting that the presence of RIα is required to facilitate this interaction. This prompted us to hypothesize that STOML2 may bind to the PKA holoenzyme, which contains both PKA Cα and RIα subunits. Structural analysis further revealed that STOML2 contains an N-terminal prohibitin homology (PHB) functional domain, which is conserved among a family of scaffold proteins (**Fig S7A**). Intriguingly, AlphaFold modeling predicted a high-confidence interaction interface between the N-terminal hairpin region of the PHB domain of STOML2 and the dimerization/docking domain of PKA RIα (**Fig. S7B**). To validate this interaction in macrophages, we used immobilized cAMP analogs to isolate either PKA regulatory subunit (8AHA-cAMP-agarose) or PKA holoenzyme (Rp-8AHA-cAMP-agarose) from macrophage lysates and found that endogenous STOML2 indeed co-precipitate efficiently with the endogenous PKA holoenzyme **(Fig. 6B).** Although PGE_2_ had no effect on STOML2 transcriptional expression, it markedly enhanced STOML2 stabilization **(Fig. 6C-D)**. Since STOML2 was reported to act as a positive regulator of PINK1 stabilization and hence is required for the efficient induction of PINK1-mediated mitophagy^38^, we next asked whether STOML2 similarly mediates PINK1 stabilization during PGE_2_-induced mitophagy in macrophages. Upon PGE_2_ stimulation, macrophages displayed robust PINK1 stabilization and Ubiquitin phosphorylation; however, both responses were substantially attenuated when STOML2 levels were depleted **(Fig. 6E-G)**, supporting that STOML2 is required for the activation of PINK1-mediated mitophagy downstream of PGE_2_-PKA signaling. Furthermore, STOML2 knockdown fully abrogated the inhibitory effects of PGE_2_ on cytosolic mtDNA levels **(Fig. 6H)** and the antiviral response **(Fig. 6I-J)**, further establishing STOML2 as a critical downstream effector and molecular hub that may connect mitophagy to regulation of STING-mediated innate immunity upon PGE_2_-PKA activation following HSV-1 infection.

**Figure 6.**
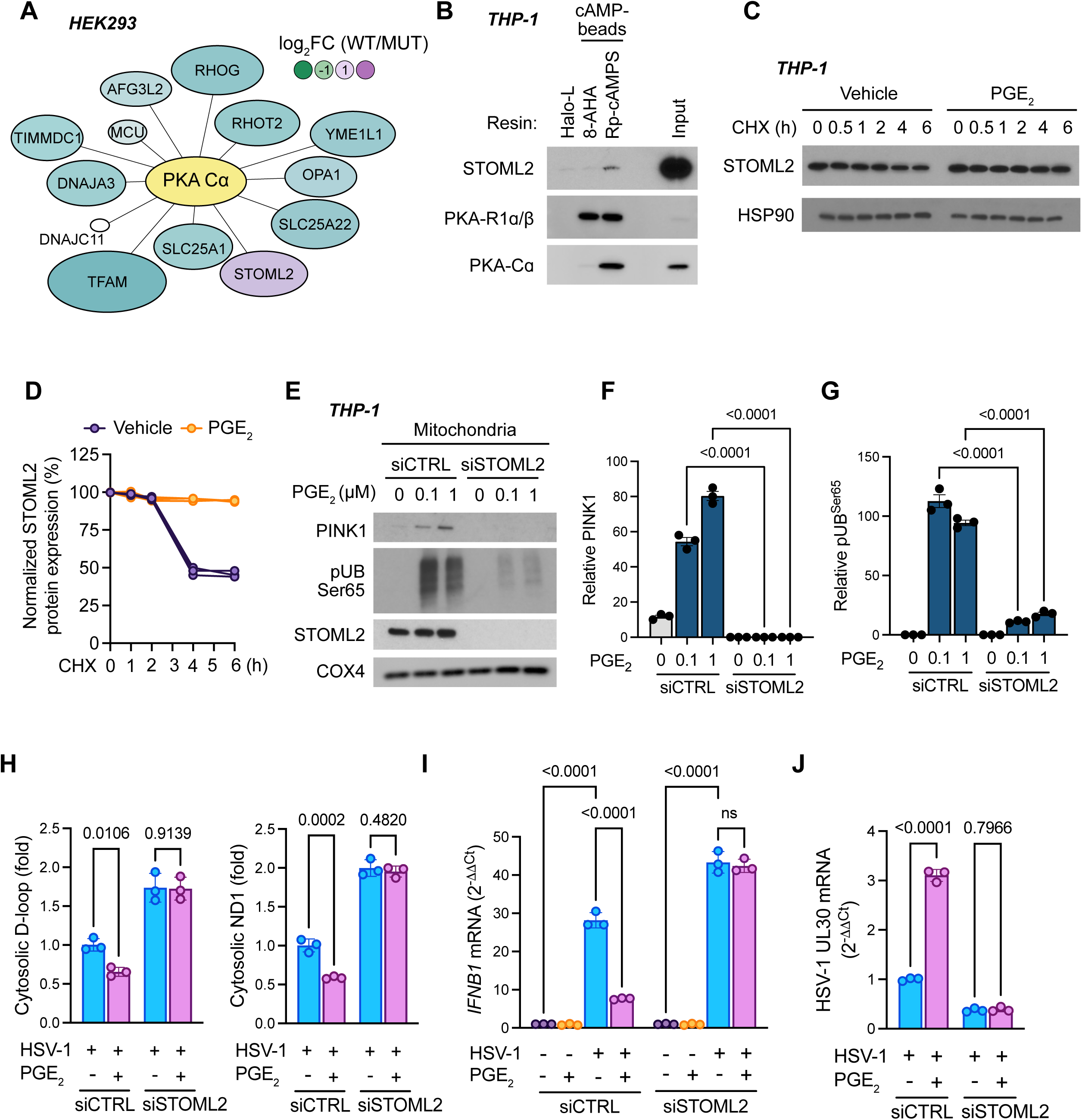
The expression of STOML2, a PKA interactor, is necessary for PINK1-mediated mitophagy induction and STING-mediated antiviral type I IFN upon PGE_2_ stimulation. **(A)** Network of mitochondrial quality control proteins interacting with wild-type (WT) and mutant (MUT) PKA Cα with BFDR σ; 0.2. Interactors are colored by log_2_ fold change (WT/MUT spectral counts) with WT-specific interactors in dark purple and MUT-specific interactors in dark green. Edges to the central bait (yellow) represent interactions detected in this study. **(B)** Whole cell lysates of THP-1 macrophages were subjected to pulldown using 8-AHA-cAMP (RIα), Rp-8-AHA-cAMPS (holoenzyme), or HaloLink resin (control), followed by immunoblotting with indicated antibodies. **(C-D)** THP-1 macrophages were stimulated with 1 µM PGE_2_ for 6 hours, followed by incubation with 1 µM cycloheximide for the indicated times. Cell lysates were collected and subjected to immunoblotting with the indicated antibodies. Representative blots from three independent experiments are shown **(C)**. Band intensities for STOML2 were quantified and normalized to HSP90 intensity (n = 3) **(D)**. **(E-G)** THP-1 macrophages were transfected with either scramble siRNA (siCTRL) or STOML2 siRNA (siSTOML2), followed by treatment with PGE_2_ at indicated concentrations for 16 hours. Mitochondrial fractions were isolated and subjected to immunoblotting with the indicated antibodies **(E)**. Band intensities for PINK1 **(F)** and pUB Ser65 **(G)** were quantified and normalized to COX4 intensity (n = 3). **(H)** THP-1 macrophages were transfected with either scramble siRNA (siCTRL) or STOML2 siRNA (siSTOML2), followed by mock or HSV-1 infection in the presence or absence of 1 µM PGE_2_. At 16 h.p.i, cytosol fractions were isolated and subjected to qPCR to assess the presence of mt-Dloop and mt-ND1 regions (n=3). **(I)** THP-1 macrophages were transfected with either scramble siRNA (siCTRL) or STOML2 siRNA (siSTOML2), followed by mock or HSV-1 infection in the presence or absence of 1 µM PGE_2_. At 16 h.p.i, whole cell lysates were collected and subjected to RT-qPCR to assess mRNA levels of IFNβ (n=3). **(J)** THP-1 macrophages were transfected with either scramble siRNA (siCTRL) or STOML2 siRNA (siSTOML2), followed by mock or HSV-1 infection in the presence or absence of 1 µM PGE_2_. At 16 h.p.i, whole cell lysates were collected and subjected to qPCR to assess HSV-1 UL30 genomic abundance (n=3). All experiments were performed with three independent biological replicates and repeated at least twice with reproducible results. Data are presented as mean ± s.e.m. Statistical significance was determined by one-way ANOVA followed by Sidak’s multiple comparisons test. *p*-values are indicated.

### PKA promotes STOML2 phosphorylation at Ser29, which is essential for PINK1-mediated mitophagy induction in response to PGE_2_

Given that PKA functions as a kinase, we next asked whether PKA modulates STOML2 phosphorylation. To examine this possibility, we performed pulldowns of a STOML2-Flag construct from HEK293 cells stimulated with forskolin and IBMX, followed by immunodetection using PKA-phosphosubstrate antibodies (PKAS, RRXX*S/T). Our data revealed that this stimulation led to a time-dependent increase in phosphorylation signal, suggesting that STOML2 may contain a consensus PKA phosphorylation motif **(Fig. 7A)**.

**Figure 7.**
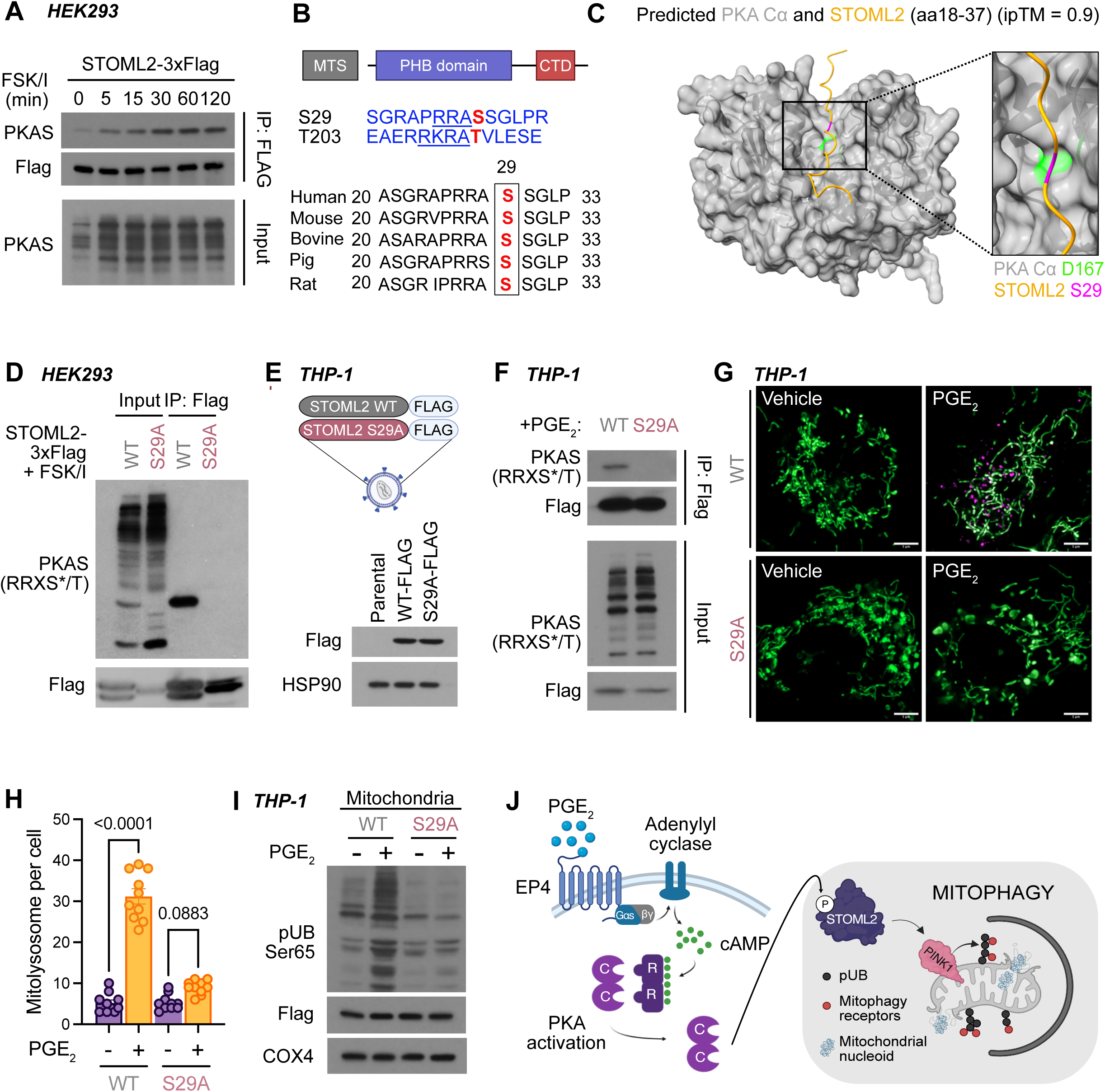
PKA promotes STOML2 phosphorylation at Ser29, which is essential for PINK1-mediated mitophagy induction in response to PGE_2_. **(A)** HEK293 cells expressing STOML2-3xFlag were treated with 1 µM forskolin and 50 µM IBMX for the indicated times. Whole cell lysates were subjected to immunoprecipitation using DYKDDDDK Fab-Trap agarose and immunoblotting with the indicated antibodies. **(B)** Top: domain architecture of STOML2 and predicted PKA phosphorylation sites (Scansite 4.0; PhosphoSitePlus). Bottom: ClustalW alignment showing evolutionary conservation surrounding STOML2 Ser29. **(C)** Representative structural model of the interaction between the active pocket of PKA Cα and STOML2 N-terminal (aa 18 to 37) (ipTM = 0.9). Gray: PKA Cα, green: PKA Cα residue D167, yellow: STOML2, magenta: STOML2 residue S29. **(D)** HEK293 cells expressing either STOML2 wild-type (WT) or S29A mutant (MUT) were treated with 1 µM forskolin and 50 µM IBMX for 6 hours. Cell lysates were subjected to immunoprecipitation using DYKDDDDK Fab-Trap agarose and immunoblotting with the indicated antibodies. **(E)** Schematic illustrating the generation of THP-1 macrophages expressing either STOML2 wild-type (WT) or S29A mutant (MUT) via lentivirus transduction. **(F)** THP-1 macrophages expressing either STOML2 wild-type (WT) or S29A mutant (MUT) were stimulated with 1 µM PGE_2_ for 16 hours. Cell lysates were subjected to immunoprecipitation using DYKDDDDK Fab-Trap agarose and immunoblotting with the indicated antibodies. **(G)** Representative Airyscan live-cell imaging of THP-1 macrophages expressing either STOML2 wild-type (WT) or S29A mutant (MUT) together with the mt-mKeima sensor were stimulated with 1 µM PGE_2_ for 16 hours. Scale bar, 10 µm. **(H)** Quantification of mitolysosome numbers from **(G)** (n = 10). **(I)** THP-1 macrophages expressing either STOML2 wild-type (WT) or S29A mutant (MUT) were stimulated with 1 µM PGE_2_ for 16 hours. Mitochondrial fractions were collected and subjected to immunoblotting with indicated antibodies. **(J)** A schematic illustrating that PKA activation promotes STOML2 phosphorylation at Ser29 which subsequently induces PINK1-mediated mitophagy in response to PGE_2_ stimulation. All experiments were performed with three independent biological replicates and repeated at least twice with reproducible results. Data are presented as mean ± s.e.m. Statistical significance was determined by one-way ANOVA followed by Sidak’s multiple comparisons test. *p*-values are indicated.

We then conducted an *in silico* analysis of potential PKA phosphorylation sites for STOML2 using the ScanSite 4.0 platform and the PhosphoSitePlus database, and identified two potential phosphorylation sites at Ser29 and Thr203 **(Fig. 7B)**. Intriguingly, AlphaFold modeling predicted that residue Ser29, but not Thr203, is in close proximity to Asp167 within the catalytic pocket of PKA Cα, a residue essential for the kinase activity of PKA Cα^31^ **(Fig. 7C)**. To assess whether PKA activation promotes STOML2 phosphorylation at Ser29, we expressed either STOML2 wild-type or S29A mutant in HEK293, followed by forskolin and IBMX stimulation. Cell lysates were then subjected to anti-Flag immunoprecipitation and PKAS immunoblotting. Strikingly, in contrast to the expression of STOML2 wild-type, expression of STOML2 S29A mutant completely abolished PKAS signal **(Fig. 7D).** To further confirm this result, we generated macrophages stably expressing either wild-type or S29A mutant STOML2 via lentiviral transduction **(Fig. 7E)**. Mimicking our findings in HEK293 cells, PGE_2_ stimulation induced robust phosphorylation of wild-type STOML2 in macrophages, whereas phosphorylation was absent in cells expressing the S29A mutant **(Fig. 7F)**. Based on these data, we concluded that PKA activation promotes STOML2 phosphorylation at Ser29.

To confirm that STOML2 phosphorylation is responsible for PGE_2_-mediated mitophagy induction, we introduced the mtKeima sensor into macrophages expressing either STOML2 wild-type or mutant. Notably, Airyscan imaging analysis revealed that the expression of the STOML2 mutant indeed fully reverted the effect of PGE_2_ on activating mitophagy **(Fig. 7G-H)**, concomitantly with an inhibition of PINK1-induced Ubiquitin phosphorylation **(Fig. 7I)**. These findings support the conclusion that STOML2 phosphorylation functions as a critical molecular switch upon PGE_2_-PKA activation to induce PINK1-mediated mitophagy which in turns removes defective mitochondria and curtail mtDNA release, thereby attenuating subsequent mtDNA-STING-mediated type I IFN and antiviral innate immunity following HSV-1 infection (**Fig. 7J**).

## DISCUSSION

Innate immunity constitutes the first line of defense against pathogen invasion and cancer development^39^. Over the last decade, the cGAS/STING pathway has emerged as a central and broadly engaged innate immune signaling pathway that is essential for robust antiviral and anticancer responses^4^. However, the cellular mechanisms that enable evasion of STING-mediated innate immunity remain elusive. Using a well-established viral infection model of STING-mediated antiviral innate immunity, we demonstrate that the most widely recognized inflammatory mediator COX2/PGE_2_ is upregulated following HSV-1 infection in an autocrine manner. This establishes a regulatory feedback loop that restrains STING-mediated type I IFN and the antiviral innate immunity to HSV-1 infection, thereby promoting viral replication.

Mitochondria have emerged as critical triggers of innate immune activation. Many viruses, including both DNA and RNA viruses, have been reported to encode proteins that, once expressed, target and damage host cell mitochondria, leading to the release of mtDNA to the cytosol and the subsequent activation of STING-mediated type I IFN and antiviral innate immunity^40–43^. In support of these findings, our results showed that HSV-1 induces mtDNA leakage into the cytosol, which, in turn, activates STING signaling and downstream type I IFN response to elicit antiviral immunity against HSV-1 infection. We unexpectedly discovered that PGE_2_ preferentially suppresses STING-mediated type I IFN and antiviral innate immunity arising from immunostimulatory cytosolic mtDNA, rather than viral DNA, generated during HSV-1 infection.

There is a fast-growing list of pathways that maintain mitochondrial homeostasis, including mitochondrial-derived vesicles (MDVs) and selective turnover of mitochondrial substructures such as nucleoids and MICOS complexes^44–47^. Among these, mitophagy, the removal of damaged or defective mitochondria by the autophagy machinery, is one of the best-characterized mechanisms that contribute to mitochondrial quality control. This can be mediated by the PTEN-induced kinase (PINK1)/Parkin pathway, mitophagy receptors, or changes in mitochondrial lipid composition^48–50^. Our study revealed that the inflammatory mediator PGE_2_ functions as an upstream physiological signal that coordinates mitochondrial quality control with the modulation of innate immunity. Mechanistically, we demonstrated that PGE_2_, via the EP4-PKA axis, robustly induces mitophagy to remove defective mitochondria and hence prevent the accumulation of immunostimulatory cytosolic mtDNA, a key trigger of cGAS activation. Intriguingly, we also found that PGE_2_ treatment leads to an increase in transcriptional expression of PGC-1α, a master activator of mitochondrial biogenesis. This suggests that mitochondrial biogenesis may be concomitantly upregulated upon mitophagy induction to replenish mitochondrial content. Given that excessive mitophagy can be detrimental due to mitochondrial depletion, we speculate that long-term suppression of STING-mediated inflammation may require coordinated engagement of both arms of mitochondrial quality control: the removal of damaged mitochondria and the production of new mitochondrial biomass.

In summary, our study supports a novel model in which COX2/PGE_2_/PKA axis functions as a negative regulator of mtDNA-STING signaling, suggesting that the inhibition of this pathway may potentiate STING-mediated type I IFN and innate immunity. Importantly, our findings establish a conceptual framework for investigating the regulatory role of COX2/PGE_2_/PKA on a myriad of physiological contexts in which mtDNA-STING signaling is implicated, including cancer, aging, and neurodegeneration.

## METHODS

### Cell culture

THP-1 (ATCC, TIB-202) was maintained in RPMI-1640 (Gibco 11875093). HEK293 (CRL-3216) (ATCC) was maintained in Dulbecco’s Modified Eagle’s Medium (DMEM, Sigma D6429). Both media were supplemented with 10% heat-inactivated fetal bovine serum (FBS) and 1X Anti-Anti (Gibco 15240062). THP-1 medium was further supplemented with 0.05 mM 2-mercaptoethanol (21-985-023). For macrophage differentiation, THP-1 cells were seeded at 5×10^5^ cells/well in 6-well plates and cultured for 72 hours with 50 ng/mL phorbol 12-myristate 13-acetate (PMA).

### Culture of bone marrow-derived macrophages

Primary murine bone marrow-derived macrophages were isolated from C57BL/6 mice and cultured in Advanced DMEM/F12 medium (Gibco, 12634010) supplemented with 10% heat-inactivated FBS, 1X Antibiotic-Antimycotic, and 20 ng/mL recombinant mouse M-CSF (PeproTech 315-02) at 37°C in a humidified 5% CO2 incubator. Bone marrow cells were cultured for 7 days, with fresh M-CSF added every 2 days. On day 7, cells were detached using trypsin-EDTA (Gibco 25300054) and seeded at 5×10^5^ cells/well in 6-well plates for experiments. Animal studies were approved by the University of California, San Diego (UCSD) Institutional Animal Care and Use Committee (IACUC) under animal study protocol (ASP) S15195 and adhered to relevant ethical regulations for animal research.

### Antibodies

Antibodies for PKA substrates (#9624), PKA Cα (#5842), PKA RIα/β (#3927), GAPDH (#2118), STING (#13647), pTBK1 S172 (5483), TBK1 (#3504), pIRF3 S396 (#4947), IRF3 (#4302), TFAM (#8076), HSP90 (#4877), pUB S65 (#62802), COX4 (#4850), pPINK1 S228 (#89010), STOML2 (#89199), Vinculin (#13901) were purchased from Cell Signaling Technology. Antibody for PINK1 (BC100-494) was purchased from Novus Biologicals, and Flag-M2 (F3165) from Millipore-Sigma. Secondary antibodies for anti-rabbit (#4030-05) and anti-mouse (#1030-05) were purchased from Southern Biotech. A complete list of primers is provided in Table S1.

### Plasmids, siRNAs and transfection

pLenti-mt-mKeima was subcloned from pCHAC-mt-mKeima (Addgene plasmid, #72342). pEYFP-N1-PINK1 plasmid was purchased from Addgene (#101874). Constructs of STOML2 wild-type and S29A mutant were purchased from VectorBuilder. cGAS and STING plasmids were generous gift from Dr. Jin Zhang (UC San Diego). To generate PKA Cα constructs, cDNAs for wild-type and mutant murine PKA Cα were transferred into the pDONR221 (12536017, Thermo Fisher) backbone using the Gateway system. Briefly, cDNAs were PCR amplified and recombined into the entry backbone using BP Clonase II according to manufacturer’s instructions. After confirming proper gene insertion with diagnostic digests, pDONR221 constructs were then transferred to the final pLVX TetOn 3XFLAG puro destination vector with the LR clonase (11791100, Invitrogen). PKA Cα constructs were tagged with a C-terminal 3xFLAG. All constructs were confirmed with sequence and functional validation.

Cells were transfected with 40 nM of SMARTpool siRNAs (Horizon Discovery) using Lipofectamine RNAiMAX reagent (#13778150, Invitrogen) according to the manufacturer’s instructions. Culture media was refreshed at 24 h after transfection. Cells were placed under serum-free conditions at 48 h and collected for experimentation at 72 h post-transfection.

### Luciferase reporter assays

THP1-ISRE-Luc2 (TIB-202-IRSE-LUC2) macrophages were seeded in 12-well plates (5×10^5^ cells/well) and treated as indicated in the figure legends. Luciferase assays were performed with the Firefly Luciferase Assay System (Promega, E1500) according to the manufacturer’s instructions.

### RNA extraction, cDNA synthesis and quantitative real-time PCR (qPCR)

THP-1 macrophages, HEK293T cells, or bone marrow-derived macrophages (BMDMs) were seeded in 6-well plates (5×10^5^ cells/well) and treated as indicated in the figure legends. Total RNA was isolated from cells using the RNeasy Mini Kit (QIAGEN, 74104) according to the manufacturer’s instructions. RNA concentration was quantified using a NanoDrop ND-1000 spectrophotometer (ThermoScientific). cDNA was synthesized from 0.5-1 μg of RNA using SuperScript VILO Master Mix (Invitrogen, 11755050) following the manufacturer’s protocol. Quantitative PCR (qPCR) was performed using Fast SYBR Green Master Mix (Applied Biosystems, 4385612) on a CFX Opus 384 Real-Time PCR system (BioRad). Three technical replicates were performed for each biological sample, and expression values for each replicate were normalized against GAPDH cDNA using the 2^-1ΔΔCt^ method. For relative expression (fold), control samples were centered at 1. A complete list of primers is provided in Table S2.

### cAMP-pulldown assay

THP-1 cells were seeded in two 10-cm plates (5×10^6^ cells per plate), differentiated into macrophages with PMA for 24 hours, and serum-starved for 12 hours. Cells were washed once with cold PBS and lysed with 1 mL of IP buffer supplemented with protease and phosphatase inhibitor cocktail, sodium orthovanadate, and IBMX. Cell extracts were scraped, collected in 1.5 mL tubes, and centrifuged at 16,000x g for 10 minutes at 4°C. Supernatants were collected, pooled, divided into equal volumes, and incubated with 20μL of 8-AHA-cAMP- or Rp-8-AHA-cAMPS-agarose (Biolog, A012 and A028) or HaloLink-agarose (Promega, G1914) as a negative binding control, with continuous rotation at 4°C for 3 hours. A 75 μL aliquot of the pooled sample was prepared with 4X Laemmli buffer as input. After incubation, agarose beads were washed three times with IP buffer, eluted with 50 μL of 2X Laemmli buffer, and heated at 95°C for 5 minutes. Pulldown and input samples were analyzed by immunoblot.

### Lentivirus production and transduction

HEK293T cells, seeded in poly-D-lysine-coated 10-cm plates, were co-transfected with pCMV-VSV (envelope vector), psPAX2 (packaging vector), and pLentiCRISPRv2 plasmids using Turbofect reagent. Supernatants were collected at 48- and 72-hours post-transfection, and lentivirus was concentrated using Lenti-X concentrator (Takara, 631232). HEK293T cells, seeded in 6-well plates, were transduced with lentiviral particles at a multiplicity of infection (MOI) of 1:1 and selected with 1 μg/mL puromycin (Gibco, A1113803) for 7 days.

### Cell stimulation and lysate preparation

THP-1 or bone marrow-derived macrophages (BMDMs) were seeded in 6-well plates (5×10^5^ cells/well) and serum-starved for 12 hours. Cells were then stimulated with 16,16-dimethyl-PGE_2_ (Cayman, 14750), forskolin (Sigma, F6886), IBMX (Sigma, I5879). For inhibition studies, cells were pre-incubated with EP4 antagonist ONO AE3-208 (Tocris 3565), PKA inhibitor BLU0588 (TargetMol, T60169), during the starvation period. Following stimulation, cells were washed once with cold PBS and lysed with 200μL of RIPA lysis buffer (Pierce 89900) supplemented with protease and phosphatase inhibitor cocktail (ThermoScientific, 78446) and sodium orthovanadate (NEB, P0758S). Cell extracts were scraped, collected, and centrifuged at 16,000xg for 10 minutes at 4°C. Supernatants were collected, protein concentrations were quantified, samples were prepared with 4X Laemmli buffer (Bio-Rad, 1610747), and heated at 95°C.

### Flag pull-down and immunoprecipitation

HEK293T cells, seeded in 6-well plates, transfected, serum-starved, and stimulated, were lysed with 1 mL of IP buffer (Pierce, 87788) supplemented with protease and phosphatase inhibitors. Cell extracts were scraped, collected, and centrifuged at 16,000xg for 10 minutes at 4°C. Supernatants were collected and incubated with 25μL of DYKDDDDK Fab-Trap agarose (Proteintech, ffa), with continuous rotation at 4°C for 2 hours. A 75 μL aliquot of supernatant was saved as input and prepared with 4X Laemmli buffer. Following incubation, agarose beads were washed three times with 1 mL of IP buffer, eluted with 50 μL of 2X Laemmli buffer, and heated at 95°C for 5 minutes. Pulldown and input samples were analyzed by immunoblot.

### Subcellular fractionation and detection of mtDNA in cytosolic extracts

Whole-cell extracts were collected for each sample before fractionation. Cytosol and mitochondrial fractionation were performed using the Mitochondrial Isolation Kit for Cultured Cells (Thermo Scientific, 89874) following manufacturer’s instructions. To detect cytosolic mtDNA, DNA was then isolated from these pure cytosolic fractionations using QIAQuick Nucleotide Removal Columns (QIAGEN, 28306). Quantitative PCR was performed on both whole-cell extracts and cytosolic fractions using nuclear DNA primers (18S) and mtDNA primers (Dloop and ND1)., and the C_t_ values obtained for mtDNA abundance for whole-cell extracts served as normalization controls for the mtDNA values obtained from the cytosolic fractions. This allowed effective standardization among samples and controlled for any variations in the total amount of mtDNA across experimental groups. A complete list of primers used for the detection of cytosolic mtDNA is provided in Supplemental Table 2.

### Western blotting

Protein samples prepared with Laemmli buffer were resolved on SDS / 10% polyacrylamide electrophoresis gels and transferred to polyvinylidene difluoride membranes (Millipore; IPVH304F0). Membranes were blocked and probed with appropriate primary antibodies overnight at 4°C. After incubation for 1h at room temperature with the secondary horseradish peroxidase–conjugated antibodies, reaction products were developed with Pierce ECL Western Blotting Substrate (Thermo Scientific;32106) or Immobilon ECL UltraPlus Western HRP Substrate (Millipore; WBULP).

### HSV-1 infection

THP1- or bone marrow-derived macrophages were seeded in 6-well plates (5×10^5^ cells/well) and infected in 1 mL serum-free DMEM with HSV-1 at 1 MOI. Plate was rocked every 30 minutes to ensure adequate coverage of the cells. After 1 h, the virus was removed from the cells and replaced with 1 mL fresh, complete media containing vehicle or PGE_2_. Total DNA was isolated using the DNeasy Blood & Tissue Kit (Qiagen) according to the manufacturer’s instructions. Relative HSV-1 genome abundance was determined using primers specific for nuclear 18S and HSV-1 UL30.

### IFNβ ELISA

Cells were stimulated as indicated, and supernatant was collected and centrifuged to remove cellular debris. Concentration of secreted IFNβ was measured using Human IFN-Beta TCM ELISA, with high sensitivity (PBL Assay Science, 41435) according to the manufacturer’s instructions.

### Intracellular cAMP ELISA

Cells were stimulated as indicated and cell lysates were collected. Concentration of intracellular cAMP was measured using the Enzo Direct cAMP kit (ADI-900-066A, Enzo) according to the manufacturer’s instructions.

### Extracellular PGE_2_ ELISA

Cells were stimulated as indicated, and cell lysates were collected. Concentration of extracellular PGE_2_ was measured using the Prostaglandin E_2_ ELISA Kit – monoclonal (Cayman, 514010) according to the manufacturer’s instructions.

### Oxygen consumption analysis

Cells were plated in XF96 plates (SeaHorse Biosciences) at 10,000 cells per well and treated as indicated in the figure legends. Once the samples are ready, cellular O_2_ was determined in a SeaHorse Bioscience XF96 extracellular flux analyzer according to the manufacturer’s instructions. Cells were maintained at 37°C in normal growth medium without serum.

### Airyscan live cell imaging

Cells were imaged in eight-well chamber slides (C8-1.5H-N, Cellvis). To quantify extramitochondrial DNA, cells were labelled with the following dyes: PicoGreen (1:500, P11495, Thermo Fisher), Mitotracker Deep Red (100 nM, M22426, Thermo Fisher). Cells were incubated with dyes for 30 minutes, followed by three washes with PBS. Cells were imaged in live cell medium composed of phenol-red-free DMEM (921-063-02, Thermo Fisher) supplemented with Prolong Antifade (P36975, Thermo Fisher) and 10% FBS. Samples were excited with 488 nm laser, followed by sequential excitation with a 561 nm laser. To measure mitolysosome numbers using mt-mKeima sensor, THP-1 macrophages expressing mt-mKeima were excited at 488 nm and 560 nm laser lines as described previously^51^. For all experiments, live cells were imaged with a Plan-Apochromat 63×/1.4 NA oil objective on an inverted Zeiss 880 LSM Airyscan confocal microscope with the environmental control system supplying 37 °C, 5% CO2, and humidity. Airyscan data were processed using Zen Black (Zeiss, version 2.3). For all Airyscan live cell images, single planes are shown.

### Image analysis

All image processing and analyses were performed using Fiji (ImageJ).

### Quantification of extramitochondrial mtDNA with Mitotracker Deep Red

Images were pre-processed by subtracting the background with a rolling ball radius of 25 pixels. A binary image of the Pico Green and Mitotracker Deep Red channel was generated using automatic thresholding (Otsu’s method for Pico Green, Li’s method for mitochondria). A binary image of non-mitochondrial Pico Green was generated by using the image calculator to subtract mitochondria from Pico Green, after which watershed was applied, and ROIs greater than or equal to 100 nm^2^ were identified using “Analyze particles”. ROIs were inspected and confirmed to be present outside of mitochondria.

### Quantification of mitolysosome numbers

Images were pre-processed by subtracting the background with a rolling ball radius of 25 pixels. A binary image of 488-nm (neutral pH) and 561-nm (acidic pH) channels was generated using automatic thresholding (Otsu’s method). A binary image of mitolysosome was generated by using the image calculator to subtract neutral pH from acidic pH, after which watershed was applied, and ROIs greater than or equal to 100 nm^2^ were identified using “Analyze particles”.

### 4D Lattice Light-Sheet Microscopy

Four-dimensional imaging was performed using a custom-built lattice light-sheet microscope developed by the Eric Betzig laboratory at HHMI Janelia Research Campus and UC Berkeley (https://www.biorxiv.org/content/10.1101/2025.06.02.657494v1.full). The system was equipped with a 0.6 NA excitation objective (Thorlabs TL20X-MPL), and a 1.0 NA detection objective (Zeiss W Plan-Apochromat 20×/1.0, model #421452-9800) and a Hamamatsu ORCA-Fusion BT sCMOS camera.

Excitation was provided by 488-nm and 560-nm lasers to image the pH-sensitive mt-Keima reporter in healthy mitochondria and in mitochondria undergoing mitophagy, respectively. Illumination was generated using a multiple Bessel-beam lattice light-sheet pattern with a numerical aperture range of NA_min = 0.35 to NA_max = 0.40. During imaging, samples were maintained at 37 °C in a humidified atmosphere containing 5% CO2. Raw lattice light-sheet microscopy data were preprocessed using the LiveLattice pipeline (https://onlinelibrary.wiley.com/doi/10.1111/jmi.13358), including camera background subtraction, photobleaching correction, sample-scan deconvolution, deskewing, and rotation into a standard Cartesian coordinate system. Each preprocessed 3D volume comprised 976 × 1536 × 270 voxels (x, y, z) with an isotropic voxel size of 0.111 µm. Time-lapse imaging was performed at 11.3-s intervals for a total of 60 frames.

### 4D Data Analysis

For each cell of interest in the 4D movies, we manually cropped the data to extract single-cell movies. To perform single-particle tracking on the mitophagy mitochondria, we used the STracking python library (https://academic.oup.com/bioinformatics/article/38/14/3671/6594114). We used the differential of Gaussian filter to segment mitophagy puncta in the 560nm excitation channel in 3D for each time point and used Euclidean distance cost to track particles across the full 60-frame movie. We then computed the features using custom-written Python analysis scripts.

### RNA-sequencing (RNA-seq) and bioinformatics analysis

Total RNA extracted from THP1 was subjected to paired-end 150bp sequencing on the Illumina NovaSeq6000 sequencer at Novogene Co., Ltd. Sequencing reads were aligned to GRCh38 human reference genome by STAR v2.7.9 using Gencode v38 annotations. Read count data was generated by RSEM, and differentially expressed genes were obtained by DESeq2^52^. Gene set enrichment analysis (GSEA) was performed using GSEA MSigDB v2023.2^53^. Transcription factor (TF) activities in the RNA-seq were inferred using R package decoupleR v2.9.1^54^. The size factor normalized gene expression matrix and stat values output from DESeq2 were used for TF activity prediction with *run_ulm* function. ISG analysis and interferon type classification were conducted using Interferome 2.0^55^. Heatmaps were generated using the tidyverse, ggplot2, viridis, RColorBrewer and pheatmap packages in R.

### Proteomic sample preparation

Affinity-purification was performed at 4°C unless otherwise stated. To begin, 500 µL of lysis buffer consisting of 50 mM Tris (pH 7.4 at 4°C), 150mM NaCl, 1mM EDTA, 0.5% NP40, 1x protease inhibitor cocktail (Roche, complete mini EDTA free), 125 U Benzonase/mL (E1014-25KU, Sigma-Aldrich) was added to each cell pellet. Lysis was achieved by two cycles of 5-10min incubation on dry ice, followed by incubation at 37°C in a water bath until all thawed. The lysate was then clarified by centrifugation at 13,000 x g for 15 min, 4°C. The clarified lysate (supernatant) was then transferred to a 96-well deep-well plate and placed on a KingFisher Flex. To each well was added 30 µL of M2 anti-FLAG beads (M8823-5ML, Sigma-Aldrich) that had been washed twice in wash buffer 2 (WB2: 50 mM Tris (pH 7.4 at 4°C), 150 mM NaCl, 1 mM EDTA. The sample was incubated with the anti-FLAG beads for 90min at 4°C with agitation. Next, the beads were washed twice with 1mL of wash buffer 1 (WB1: 50 mM Tris (pH 7.4 at 4°C), 150 mM NaCl, 1 mM EDTA, 0.05% NP40), followed by two additional washes of 1mL of WB2. Finally, enriched proteins were then eluted by the addition of 30µL of FLAG elution buffer (100 µg/ml 3xFLAG peptide (F3290-4MG, Sigma-Aldrich) in 0.05% RapiGest (186001861, Waters Corporation), 50 mM Tris (pH 7.4 at 4°C), 150 mM NaCl, 1 mM EDTA) for 15min at room temperature with manual agitation. A magnetic plate was then used to aggregate the beads, and the supernatant was transferred to a new 96-well plate. This elution from the beads using the FLAG elution buffer was repeated, and the resulting supernatant was added to the initial elution. Next, 15µL of 8M urea, 250mM Tris, 5mM DTT were added to each sample, followed by incubation at 60°C for 15min. The samples were then cooled to room temperature, followed by the addition of iodoacetamide to a final concentration of 3 mM, and incubated for 45 min at room temperature in the dark. Next, 3 mM dithiothreitol and 1 µg of trypsin (V5113, Promega) were added to the samples, which were incubated at 37°C overnight. The next day, samples were acidified with 10% TFA to a final pH below 2, then incubated at room temperature for 30 min. Lastly, the samples were desalted on C18 tips (HNS S18V, NEST Group), lyophilized to dryness, and resuspended in 15µL of 4% formic acid, 2% acetonitrile.

### Mass spectrometry data acquisition and analysis

2µL of each sample was injected into an Orbitrap Fusion Tribrid Mass Spectrometer (Thermo Scientific) operated in positive ion mode using an EASY-nLC 1200 UHPLC (Thermo Scientific). Peptides were separated over a 75-μm internal diameter × 25-cm long picotip column packed with 1.9 μM C18 particles (Dr. Maisch) at a flow rate of 300nL/min. Buffer A was comprised of 0.1% formic acid, and buffer B was comprised of 80% acetonitrile in 0.1% formic acid. Peptides were eluted by a linear gradient of 5 to 38% B over 44 min, followed by a 5 min ramp to 100% bubber B, and a subsequent 20min wash at 100% buffer B. MS data was collected over a 70 min total acquisition time with MS1 detection in the Orbitrap at 120K resolution in profile mode, 300-1500 m/z scan range, a 100ms maximum injection time, a 2×10^5^ AGC target, a 60% RF lens setting, a 275°C transfer tube temperature, and a 2kV spray voltage. Data-dependent MS2 scans were performed in the ion trap in centroid mode on charge states from 2-6, with a +/- 10ppm exclusion for a duration of 20s after two observations. Selected precursors were isolated with a 1.6 m/z window and fragmented with a 30% normalized HCD energy in the ion trap with a rapid scan rate, a 110 m/z first mass, automated scan range selection, a 35ms maximum injection time, a 1×10^4^ AGC target.

Raw data were searched against the canonical human protein sequences from Uniprot (downloaded March 21, 2018) using Maxquant (version 1.6.6.0)^56^. Default search parameters were used, including a minimum peptide length of 7 amino acids, variable modification of acetylation of the protein N-terminus, variable modification of oxidation on methionine, match between runs was disabled, and peptides and protein were filtered to a 1% false discovery rate. Spectral count values were then analyzed by SAINTexpress^57^ to prioritize a set of high confidence interaction partners (BFDR < 0.05). Finally, MSstats^58^ was used to quantify changes in interaction partners between wild-type and mutant PKA Cα.

### Structural prediction

STOML2-PKA RIα and STOML2-PKA Cα complexes were modeled from sequences via AlphaFold3. AlphaFold3 was run through its public web server, which generated five models per complex, ranked by the AlphaFold3 ranking score. To ensure an accurate modeling of PKA Cα active state, we included as inputs two Mg^2+^ ions, one molecule of ATP, and the phosphorylation of residues T198 and S339, which are relevant PTMs for kinase activity. Twenty-five models were generated, and the models with the highest ipTM scores were selected as the representative structures. ChimeraX1.8 was used to analyze the predicted structure.

### Quantification and statistical analysis

The sample size and number of repeats are indicated in the respective figure legends. Statistical significance was calculated by one-way or two-way ANOVA (more than two groups) with Sidak comparisons or a two-tailed Student’s t test (two groups), as indicated in the figure legends. Spearman’s rank correlation p coefficient (r) was used to assess the correlation between transcript levels and individual age. No animals or data points were excluded from the analyses. All statistical tests were performed with GraphPad Prism 10.0.1 for Mac (GraphPad Software; www.graphpad.com), and data were presented as the mean ± standard error of the mean (s.e.m).

## Data availability

Bulk RNA-seq sequencing data have been deposited in Gene Expression Omnibus (GEO) with accession number GSE324662. All raw MS data files, search results and individual spectral libraries are available from the Pride partner ProteomeXchange Consortium via the PRIDE partner repository^59^ with the dataset identifier: PXD075490, and the user token: pDMIaejNPRdD.

## ACKNOWLEDGMENTS

We thank Professor Susan S. Taylor from UC San Diego for providing PKA plasmid constructs and for her perspectives and insights to this study. We thank Professor Thomas Riffelmacher from La Jolla Institute for Immunology for assistance with mitochondrial respirometry assay. We thank Professor Jin Zhang from UC San Diego for kindly providing cGAS and STING plasmid constructs. Diagrams were generated using BioRender.

Research in the J.S.G. lab is supported by grants R01DE033909, R01DE026644, and U54CA274502. Research in the N.J.K. lab is supported by the National Institute of Health (NIH) grant U54CA209891. Research in the J.S. lab is supported by the NIH Director’s New Innovator Award grant 1R01GM148765-01. Research in the U.M. lab is supported by the Chan Zuckerberg Initiative DAF (CZI Imaging Scientist Award DOI:10.37921/694870itnyzk), the Goeddel Family Technology Sandbox, the MFN2 Foundation and the Dr. David V. Goeddel Chancellor’s Endowed Chair in Biological Sciences. P.T.T.V. is supported by the Cancer Cell Map Initiative grant U54CA274502 and the Tobacco-related Disease Research Program predoctoral fellowship grant T34DT8340. T.S.H. is supported by the Tobacco-related Disease Research Program predoctoral fellowship grant T32DT4965 and the National Cancer Institute (NCI) grant T32CA121938. K.S. is supported by the Takeda Science Foundation research fellowship. Z.W. is supported by the American Heart Association Predoctoral fellowship grant 24PRE1196243.

## AUTHOR CONTRIBUTIONS

Conceptualization: P.T.T.V., S.R.A.-G., T.S.H., D.J.R., J.S.G..; methodology: P.T.T.V., J.C., Z.W., H.H., S.R.A.-G., T.S.H., D.J.R., K.S., E.S. Y.Z., K.F.; investigation: P.T.T.V., S.R.A.-G., T.H.S.; visualization: P.T.T.V., J.C., Z.W., H.H., D.J.R., Y.Z.; funding acquisition: N.J.K., J.S., U.M., J.S.G., project administration: J.S.G.; supervision: J.S.G.; writing – original draft: P.T.T.V., J.S.G.; writing – review & editing: P.T.T.V., S.R.A.-G, K.S, U.M, J.S.G. All authors read and approved the version of the manuscript.

