## Supplemental Materials for "The COX2-PGE2-PKA Axis Suppresses Antiviral Immunity by Inhibiting mtDNA-Dependent STING Activation"

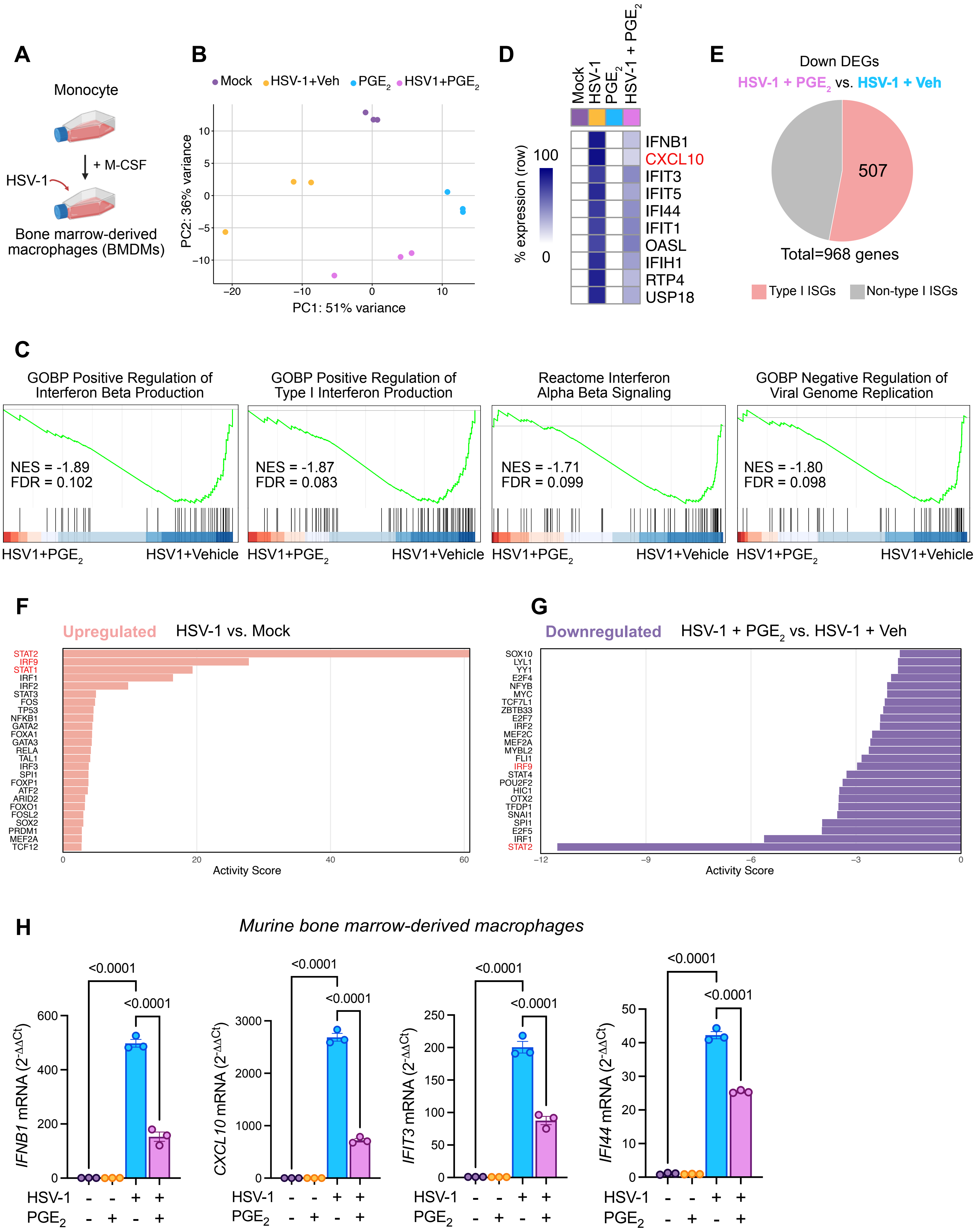

A
Monocyte
+ M-CSF
HSV-1
Bone marrow-derived macrophages (BMDMs)
B
Mock
HSV-1+Veh
PGE2
HSV1+PGE2
PC2: 36% variance
PC1: 51% variance
D
Mock
HSV-1
PGE2
HSV-1 + PGE2
% expression (row)
100
0
IFNB1
CXCL10
IFIT3
IFIT5
IFI44
IFIT1
OASL
IFIH1
RTP4
USP18
E
Down DEGs
HSV-1 + PGE2 vs. HSV-1 + Veh
507
Total=968 genes
Type I ISGs
Non-type I ISGs
C
GOBP Positive Regulation of Interferon Beta Production
NES = -1.89
FDR = 0.102
GOBP Positive Regulation of Type I Interferon Production
NES = -1.87
FDR = 0.083
Reactome Interferon Alpha Beta Signaling
NES = -1.71
FDR = 0.099
GOBP Negative Regulation of Viral Genome Replication
NES = -1.80
FDR = 0.098
HSV1+PGE2
HSV1+Vehicle
F
Upregulated
HSV-1 vs. Mock
STAT2
IRF9
STAT1
IRF1
IRF2
STAT3
FOS
TP53
NFKB1
GATA2
FOXA1
GATA3
RELA
TAL1
IRF3
SPI1
FOXP1
ATF2
ARID2
FOXO1
FOSL2
SOX2
PRDM1
MEF2A
TCF12
Activity Score
G
Downregulated
HSV-1 + PGE2 vs. HSV-1 + Veh
SOX10
LYL1
YY1
E2F4
NFYB
MYC
TCF7L1
ZBTB33
E2F7
IRF2
MEF2C
MEF2A
MYBL2
FLI1
IRF9
STAT4
POU2F2
HIC1
OTX2
TFDP1
SNAI1
SPI1
E2F5
IRF1
STAT2
Activity Score
H
Murine bone marrow-derived macrophages
IFNB1 mRNA (2^-ΔΔCt)
CXCL10 mRNA (2^-ΔΔCt)
IFIT3 mRNA (2^-ΔΔCt)
IFI44 mRNA (2^-ΔΔCt)
<0.0001
<0.0001
HSV-1 - - + +
PGE2 - + - +

### Figure S1, related to Figure 1.

**(A)** Schematic illustrating the experimental workflow for bone marrow-derived macrophages infected with HSV-1.

**(B)** Principal component analysis (PCA) of bulk RNA-seq data for the indicated groups.

**(C)** Enrichment plots for the indicated gene sets showing gene enrichment between HSV-1+ $PGE_2$ and HSV-1 + Veh macrophages. NES, normalized enrichment score; FDR, false discovery rate.

**(D)** Heatmaps for selected type I ISGs differentially expressed between the indicated groups. Regularized log2 expression values are row-mean subtracted.

**(E)** Interferome analysis of downregulated differentially expressed genes (DEGs) in HSV-1+$PGE_2$ versus HSV-1+Veh macrophages, showing the proportion of type I ISGs (52.3%).

**(F-G)** Horizontal bar plot showing the predicted transcription activities comparing HSV-1 versus mock **(F)** and HSV-1 + PGE2 versus HSV-1 + Veh **(G)** macrophages.

**(H)** Bone marrow-derived macrophages were mock-infected or infected with HSV-1 (MOI = 1) in the presence or absence of 1 μM $PGE_2$. At 16 h.p.i, cell lysates were collected and subjected to RT-qPCR to measure mRNA levels of representative type I ISGs via qRT-PCR (n = 3). Data are shown $\Delta\Delta$Ct values (mean ± s.e.m). Statistical significance was determined by one-way ANOVA followed by Sidak's multiple comparisons test; *p*-values are indicated.

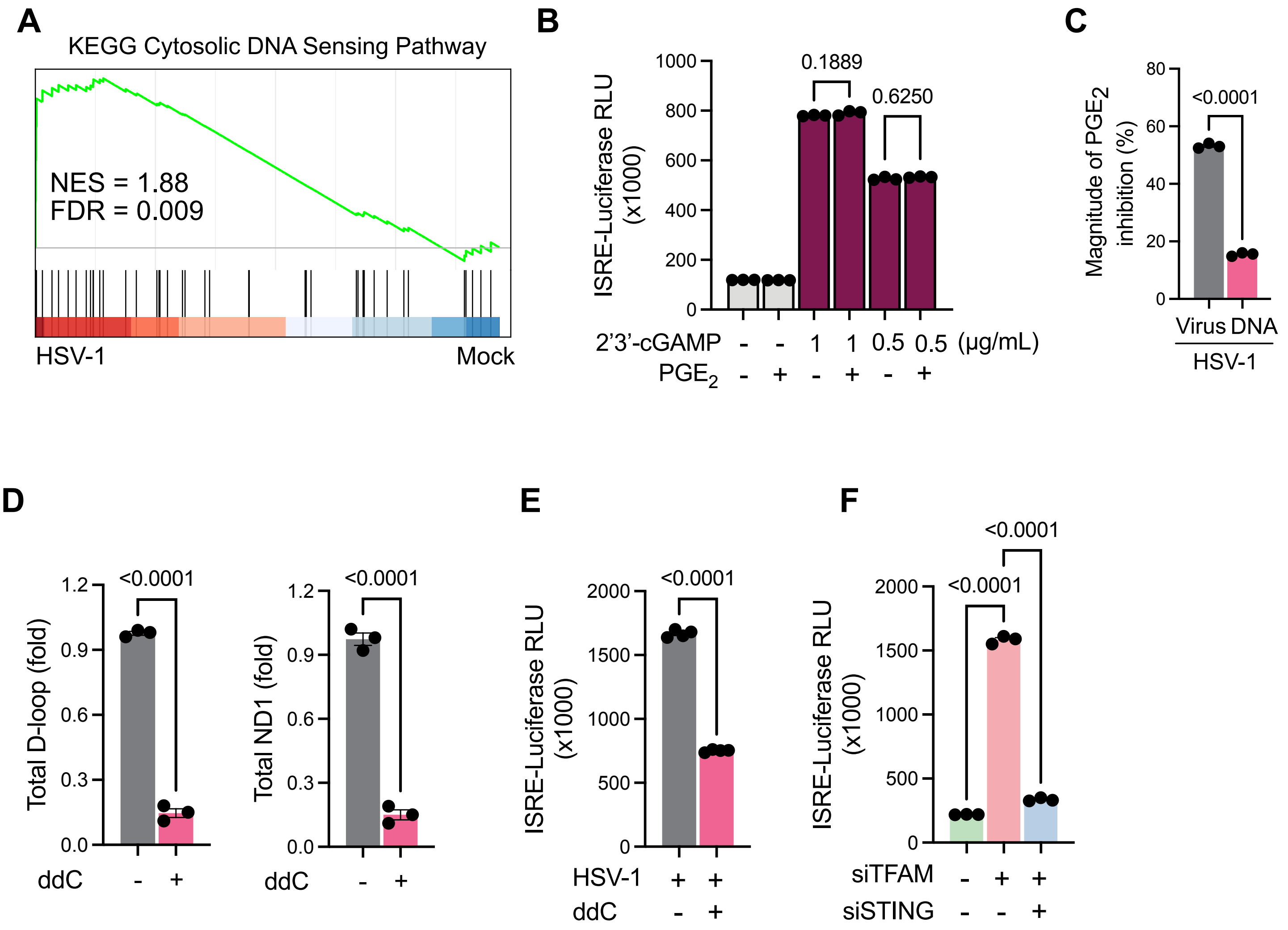

A
KEGG Cytosolic DNA Sensing Pathway
NES = 1.88
FDR = 0.009
HSV-1
Mock
B
ISRE-Luciferase RLU (x1000)
1000
800
600
400
200
0
0.1889
0.6250
2'3'-cGAMP - - 1 1 0.5 0.5 (μg/mL)
PGE2 - + - + - +
C
Magnitude of PGE2 inhibition (%)
80
60
40
20
0
<0.0001
Virus DNA
HSV-1
D
Total D-loop (fold)
1.2
0.9
0.6
0.3
0.0
<0.0001
ddC - +
Total ND1 (fold)
<0.0001
ddC - +
E
ISRE-Luciferase RLU (x1000)
2000
1500
1000
500
0
<0.0001
HSV-1 + +
ddC - +
F
ISRE-Luciferase RLU (x1000)
2000
1500
1000
500
0
<0.0001
<0.0001
siTFAM - + +
siSTING - - +

**Figure S2, related to Figure 2.**

**(A)** Enrichment plot for the indicated gene set showing gene enrichment between HSV-1+ $PGE_2$ and HSV-1 + Veh macrophages.

**(B)** THP-1 macrophages expressing ISRE-Luciferase were transfected with STING agonist, 2'3'-cGAMP, at indicated concentrations in the presence or absence of 1 µM $PGE_2$ for 16 hours. Cell lysates were collected and subjected to a luciferase reporter assay to assess ISRE promoter activity (n = 3).

**(C)** The magnitude of PGE2-mediated inhibition was calculated by comparing macrophages infected with HSV-1 (MOI = 1) to macrophages transfected with viral DNA at the indicated concentrations (n = 3).

**(D)** THP-1 macrophages were treated with control or 10 µM ddC for 16 hours. Cell lysates were collected and subjected to qPCR to assess levels of total mt-D-loop and mt-ND1 regions (n=3).

**(E)** THP-1 macrophages expressing ISRE-Luciferase were mock-infected or infected with HSV-1 (MOI = 1) in the presence or absence of 10 µM ddC. Cell lysates were collected and subjected to a luciferase reporter assay to assess ISRE promoter activity (n = 3).

**(F)** THP-1 macrophages expressing ISRE-Luciferase were transfected with either control siRNA or TFAM siRNA, in the presence or absence of STING siRNA. Cell lysates were collected and subjected to a luciferase reporter assay to assess ISRE promoter activity (n = 3).

All experiments were performed with three independent biological replicates and repeated at least twice with reproducible results. Data are presented as mean ± s.e.m. Statistical significance was determined by one-way ANOVA followed by Sidak's multiple comparisons test (**B**, **F**) or two-tailed Student's t-test (**C-E**). *p*-values are indicated.

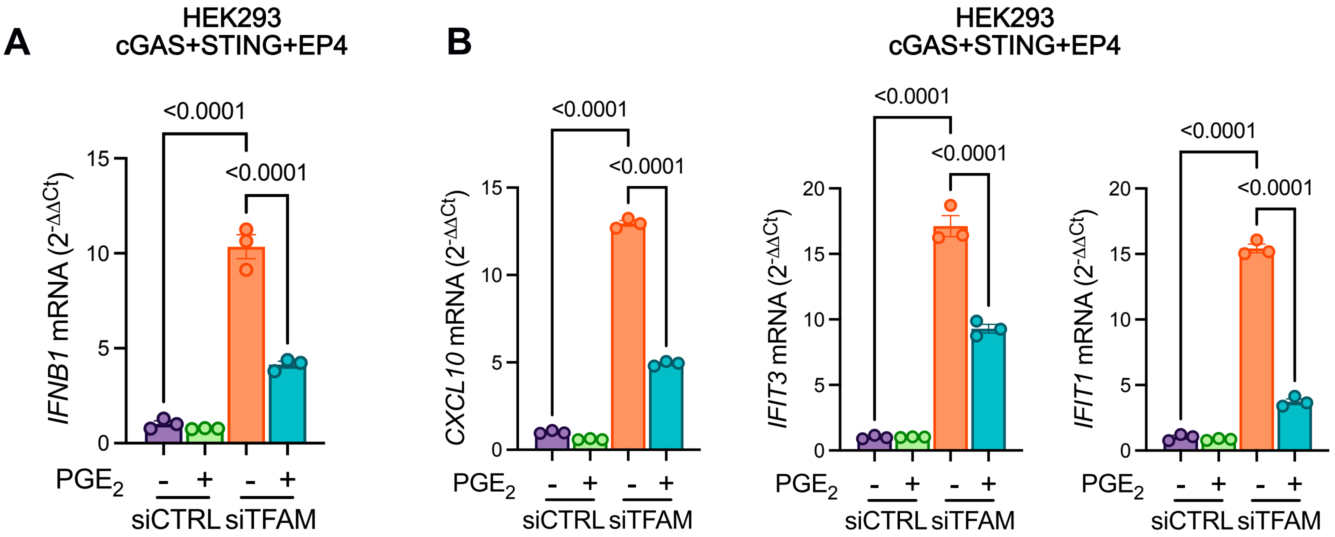

A
HEK293
cGAS+STING+EP4
<0.0001
<0.0001
IFNB1 mRNA (2-ΔΔCt)
PGE2
siCTRL
siTFAM
B
HEK293
cGAS+STING+EP4
CXCL10 mRNA (2-ΔΔCt)
IFIT3 mRNA (2-ΔΔCt)
IFIT1 mRNA (2-ΔΔCt)

**Figure S3, related to Figure 2.**

**(A-B)** HEK293 cells were transfected with either control siRNA or TFAM siRNA, followed by transfection with cGAS, STING, and EP4 plasmids. Cells are then stimulated with either vehicle or 1 μM $PGE_2$ for 16 hours. Cell lysates were collected and subjected to RT-qPCR to assess mRNA levels of IFNβ (n = 3) **(A)** and representative type I ISGs (n = 3) **(B)**.

Data are presented as mean ± s.e.m. Statistical significance was determined by one-way ANOVA followed by Sidak's multiple comparisons test. *p*-values are indicated.

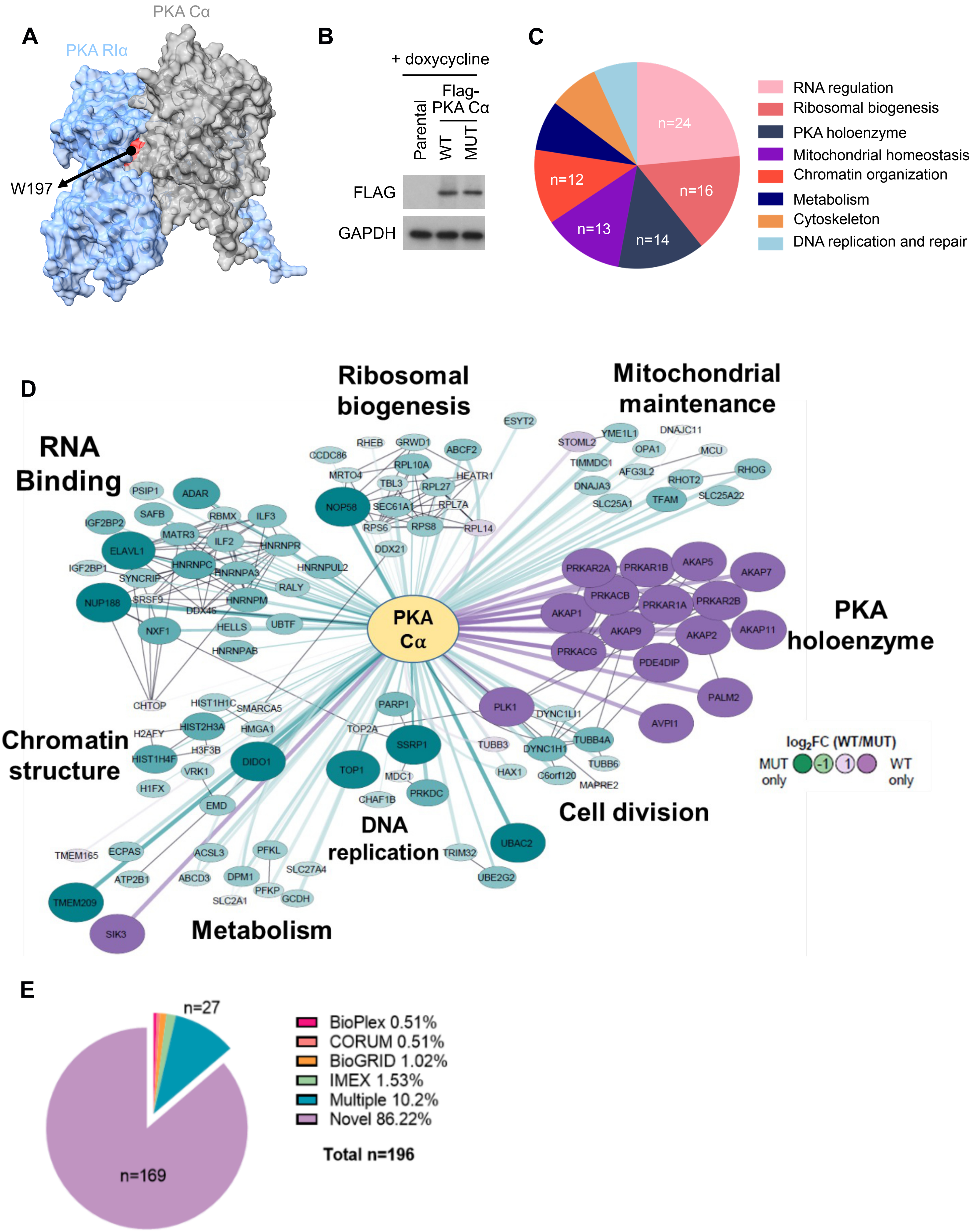
A
PKA Cα
PKA RIα
W197
B
+ doxycycline
Flag-PKA Cα
Parental
WT
MUT
FLAG
GAPDH
C
n=24
n=16
n=14
n=13
n=12
RNA regulation
Ribosomal biogenesis
PKA holoenzyme
Mitochondrial homeostasis
Chromatin organization
Metabolism
Cytoskeleton
DNA replication and repair
D
RNA Binding
Ribosomal biogenesis
Mitochondrial maintenance
PKA holoenzyme
Chromatin structure
DNA replication
Cell division
Metabolism
PKA Cα
log2FC (WT/MUT)
MUT only
-1
1
WT only
PSIP1
ADAR
SAFB
IGF2BP2
RBMX
ILF3
MATR3
ILF2
HNRNPR
ELAVL1
HNRNPC
HNRNPA3
HNRNPUL2
IGF2BP1
SYNCRIP
RALY
NUP188
SRSF9
HNRNPM
DDX46
NXF1
HELLS
UBTF
HNRNPAB
CHTOP
RHEB
GRWD1
ABCF2
CCDC86
RPL10A
MRTO4
HEATR1
TBL3
RPL27
NOP58
SEC61A1
RPL7A
RPS6
RPS8
RPL14
DDX21
ESYT2
STOML2
YME1L1
DNAJC11
OPA1
MCU
TIMMDC1
AFG3L2
RHOG
DNAJA3
RHOT2
SLC25A1
TFAM
SLC25A22
PRKAR2A
PRKAR1B
AKAP5
AKAP7
PRKACB
PRKAR1A
PRKAR2B
AKAP1
AKAP9
AKAP2
AKAP11
PRKACG
PDE4DIP
PALM2
AVPI1
PLK1
DYNC1LI1
TUBB4A
DYNC1H1
TUBB3
TUBB6
HAX1
C6orf120
MAPRE2
HIST1H1C
SMARCA5
H2AFY
HIST2H3A
HMGA1
H3F3B
HIST1H4F
DIDO1
VRK1
H1FX
EMD
PARP1
TOP2A
SSRP1
TOP1
MDC1
PRKDC
CHAF1B
UBAC2
TRIM32
UBE2G2
TMEM165
ECPAS
ACSL3
PFKL
ATP2B1
ABCD3
DPM1
SLC27A4
PFKP
GCDH
SLC2A1
TMEM209
SIK3
E
n=27
n=169
BioPlex 0.51%
CORUM 0.51%
BioGRID 1.02%
IMEX 1.53%
Multiple 10.2%
Novel 86.22%
Total n=196

**Figure S4, related to Figure 3.**

**(A)** Representative structural model of the interaction between PKA C$\alpha$ (gray) and RI$\alpha$ (cyan) with PKA C$\alpha$ residue W197 highlighted in red. The model was generated using AlphaFold 3.0.

**(B)** HEK293 parental cells harboring doxycycline-inducible Flag-tagged PKA wild-type (WT) or W197 mutant (MUT) constructs were treated with 1 µg/mL doxycycline for 48 hours. Cell lysates were collected and subjected to immunoblotting with the indicated antibodies.

**(C)** Quantification of identified protein–protein interactions (PPIs) categorized into the indicated biological functional groups. A total of 196 PPIs were analyzed (BFDR$\leq$0.2).

**(D)** PKA C$\alpha$ interaction network. All wild-type (WT) and mutant (MUT) PKA C$\alpha$ interacting prey with BFDR$\leq$0.2. Interactors are colored by $\log_2$ fold change (WT/MUT average spectral counts) with WT-specific interactors in dark purple and MUT-specific interactors in dark green. Interactors are organized by biological functional groups, and network edges between prey species reflect high-confidence interactions (STRING score >0.9). Edges connecting to the central bait (yellow) reflect interactions detected in this study.

**(E)** Quantification of novel protein-protein interactions (PPIs) against publicly available databases. The percentages of common or novel PPIs are shown. A total of 196 PPIs were considered (BFDR$\leq$0.2).

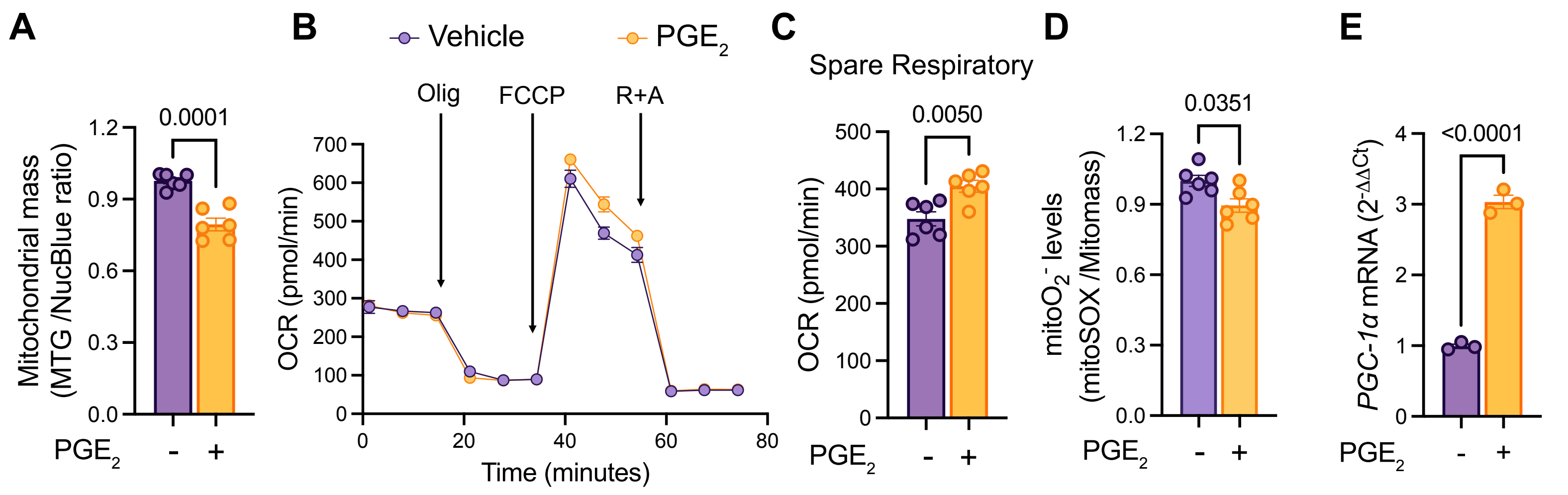

A
B
C
D
E
Vehicle
PGE2
Mitochondrial mass (MTG /NucBlue ratio)
0.0001
PGE2
-
+
Olig
FCCP
R+A
OCR (pmol/min)
Time (minutes)
Spare Respiratory
0.0050
OCR (pmol/min)
mitoO2- levels (mitoSOX /Mitomass)
0.0351
PGC-1α mRNA (2-ΔΔCt)
<0.0001

**Figure S5, related to Figure 4.**

**(A)** Mitochondrial mass was assessed using MitoTracker Green (MTG) in THP-1 macrophages treated with vehicle or 1 μM PGE2 for 24 hours (n = 5).

**(B)** THP-1 macrophages treated with vehicle or 1 μM $PGE_2$ for 24 hours were subjected to mitochondrial respirometry analysis using Seahorse Bioscience XF96 after sequential injection of Oligomycin, FCCP, and Rotenone + Antimycin (n = 6 biological replicates).

**(C)** Quantification of spare respiratory capacity derived from the Seahorse analysis in **(C)**.

**(D)** Mitochondrial superoxide production was measured as the MitoSOX Red/MTG ratio in THP-1 macrophages treated with vehicle or 1 μM $PGE_2$ for 24 hours (n = 3).

**(E)** THP-1 macrophages were treated with vehicle or 1 μM $PGE_2$ for 24 hours. Cell lysates were collected and subjected to RT-qPCR to assess mRNA levels of PGC-1$\alpha$ (n = 3).

Data are presented as mean ± s.e.m. Statistical significance was determined by a two-tailed Student's t-test. *p*-values are indicated.

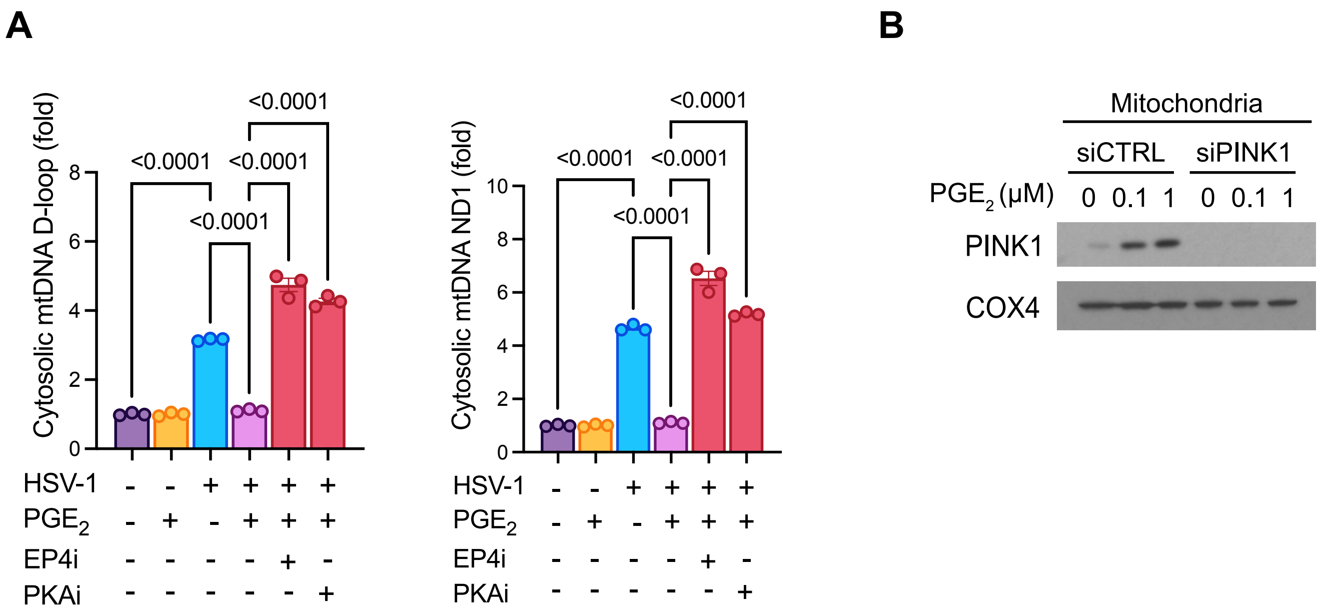
A
Cytosolic mtDNA D-loop (fold)
<0.0001
<0.0001
<0.0001
<0.0001
Cytosolic mtDNA ND1 (fold)
<0.0001
<0.0001
<0.0001
<0.0001
HSV-1
PGE2
EP4i
PKAi
B
Mitochondria
siCTRL
siPINK1
PGE2 (μM)
0 0.1 1 0 0.1 1
PINK1
COX4

**Figure S6, related to Figure 5.**

**(A)** THP-1 macrophages were infected with HSV-1 (MOI = 1), followed by 1 μM $PGE_2$ stimulation in the presence of either 1 μM EP4 inhibitor (EP4i) or 1 μM PKA inhibitor (PKAi). At 16 h.p.i, cytosol fractions were isolated and subjected to qPCR to assess levels of mt-D-loop and mt-ND1 regions (n = 3). Statistical significance was determined by one-way ANOVA followed by Sidak's multiple comparisons test. *p*-values are indicated.

(**B)** THP-1 macrophages were transfected with either scramble siRNA (siCTRL) or PINK1 siRNA (siPINK1), followed by treatment with $PGE_2$ at indicated concentrations for 16 hours. Mitochondrial fractions were isolated and subjected to immunoblotting with the indicated antibodies.

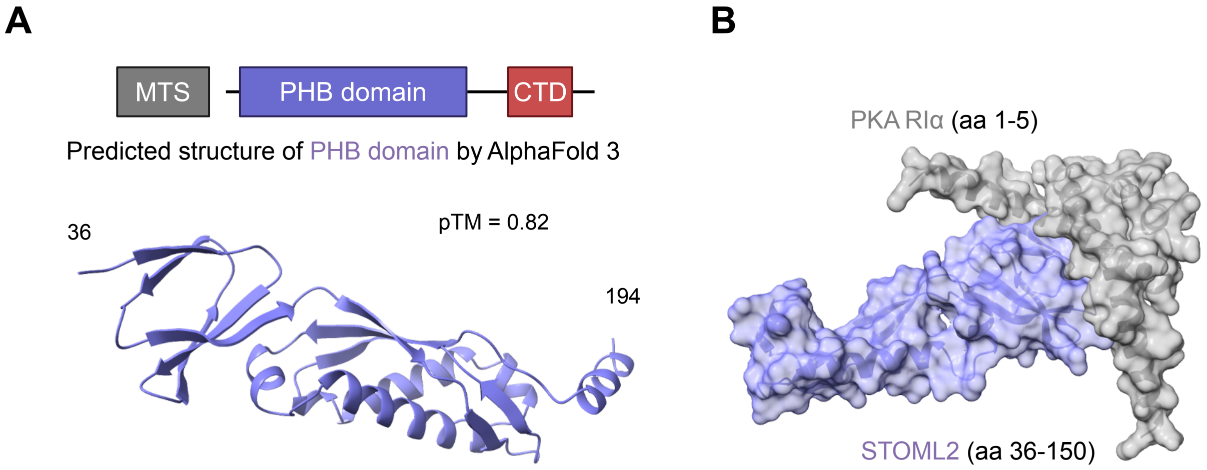
A
MTS
PHB domain
CTD
Predicted structure of PHB domain by AlphaFold 3
36
pTM = 0.82
194
B
PKA RIα (aa 1-5)
STOML2 (aa 36-150)

**Figure S7, related to Figure 6.**

**(A)** Representative structural model of STOML2 predicted using AlphaFold 3.0 (ipTM = 0.82).

**(B)** Representative structural model of the interaction between STOML2 (aa 36 to 150) and PKA Riα (aa 1 to 5) predicted using AlphaFold 3.0 (pTM = 0.81).

**Table S1. Antibodies**

| Antibody | Source | Identifier |
|---|---|---|
| Phospho-PKA substrate (RRXS*/T*) | Cell Signaling Technology | 9624 |
| PKA-Cα | Cell Signaling Technology | 5842 |
| PKA RIα/β | Cell Signaling Technology | 3927 |
| GAPDH | Cell Signaling Technology | 2118 |
| STING | Cell Signaling Technology | 13647 |
| Phospho-TBK1 S172 | Cell Signaling Technology | 5483 |
| TBK1 | Cell Signaling Technology | 3504 |
| Phospho-IRF3 S396 | Cell Signaling Technology | 4947 |
| IRF3 | Cell Signaling Technology | 4302 |
| TFAM | Cell Signaling Technology | 8076 |
| HSP90 | Cell Signaling Technology | 4877 |
| Phospho-Ubiquitin S65 | Cell Signaling Technology | 62802 |
| COX4 | Cell Signaling Technology | 4850 |
| Phospho-PINK1 S228 | Cell Signaling Technology | 89010 |
| STOML2 | Cell Signaling Technology | 89199 |
| Vinculin | Cell Signaling Technology | 13901 |
| PINK1 | Novus Biologicals | BC100-494 |
| Flag-M2 | Millipore-Sigma | F3165 |
| anti-rabbit | Southern Biotech | 4030-05 |
| Anti-mouse | Southern Biotech | 1030-05 |

**Table S2. Primers for qPCR, related to Method.**

| Identifier | | Sequence |
|---|---|---|
| Human IFNβ | Forward | TTGTTGAGAACCTCCTGGCT |
| | Reverse | TGACTATGGTCCAGGCACAG |
| Human CXCL10 | Forward | GTGGCATTCAAGGAGTACCTC |
| | Reverse | TGATGGCCTTCGATTCTGGATT |
| Human IFI44 | Forward | ATGGCAGTGACAACTCGTTTG |
| | Reverse | TCCTGGTAACTCTCTTCTGCATA |
| Human IFIT3 | Forward | TCAGAAGTCTAGTCACTTGGGG |
| | Reverse | ACACCTTCGCCCTTTCATTTC |
| Human IFIT1 | Forward | TTGATGACGATGAAATGCCTGA |
| | Reverse | CAGGTCACCAGACTCCTCAC |
| Human COX2 | Forward | CTGGCGCTCAGCCATACAG |
| | Reverse | CGCACTTATACTGGTCAAATCCC |
| HSV-1 UL30 | Forward | ATCACCGACCCGGAGAG |
| | Reverse | CAGGCGCTTGTTGGTGT |
| Human mtDNA-Dloop | Forward | CATAAAGCCTAAATAGCCCACACG |
| | Reverse | CCGTGAGTGGTTAATAGGGTGATA |
| Human mtDNA-ND1 | Forward | GAACTAGTCTCAGGCTTCAACATCG |
| | Reverse | CTAGGAAGATTGTAGTGGTGAGGGTG |
| Human 18S | Forward | TAGAGGGACAAGTGGCGTTC |
| | Reverse | CGCTGAGCCAGTCAGTG |
| Human Beta actin | Forward | ACTGGAACGGTGAAGGTGAC |
| | Reverse | AGAGAAGTGGGGTGGCTTTT |
| Human GAPDH | Forward | GGAGCGAGATCCCTCCAAAAT |
| | Reverse | GGCTGTTGTCATACTTCTCATGG |
| Human PINK1 | Forward | GCCTCATCGAGGAAAAACAGG |
| | Reverse | GTCTCGTGTCCAACGGGTC |
| Human STOML2 | Forward | AGGGGCTCTCTACTGGCTTC |
| | Reverse | ATCCGGTCTAACACAGGGATG |
| Human PGC-1α | Forward | TCTGAGTCTGTATGGAGTGACAT |
| | Reverse | CCAAGTCGTTCACATCTAGTTCA |
| Mouse Ifnβ | Forward | AGCTCCAAGAAAGGACGAACA |
| | Reverse | GCCCTGTAGGTGAGGTTGAT |
| Mouse Cxcl10 | Forward | CCAAGTGCTGCCGTCATTTTC |
| | Reverse | GGCTCGCAGGGATGATTTCAA |
| Mouse Ifit3 | Forward | CCTACATAAAGCACCTAGATGGC |
| | Reverse | ATGTGATAGTAGATCCAGGCGT |
| Mouse Ifit1 | Forward | GCCTATCGCCAAGATTTAGATGA |
| | Reverse | TTCTGGATTTAACCGGACAGC |
| Mouse Gapdh | Forward | AGGTCGGTGTGAACGGATTTG |
| | Reverse | TGTAGACCATGTAGTTGAGGTCA |
| Mouse Beta actin | Forward | GGCTGTATTCCCCTCCATCG |
| | Reverse | CCAGTTGGTAACAATGCCATGT |
